# Saturation Genome Editing reveals the functional impact of RAD51D *and* XRCC2 variants

**DOI:** 10.64898/2026.06.12.731983

**Authors:** Silvia Casadei, Matthew W. Snyder, Ivan Woo, Nahum Smith, Sabrina Best, Riddhiman K. Garge, Leslie Rodriguez-Salas, Cameron Wenman, Obsa Seid, Airi Hosokai, Alicia Xu, Malvika Tejura, Pankhuri Gupta, Sarah Heidl, Lara Muffley, Jane Ranchalis, Ross Stewart, Noah J. Goff, Shelby L. Hemker, John Baierl, Ed M. Dicks, Paul Pharoah, Lene Bjornkjaer, Karina Roenlund, Jacob O. Kitzman, Kara A. Bernstein, Andrew B. Stergachis, Predrag Radivojac, Douglas M. Fowler, Lea M. Starita

## Abstract

Germline pathogenic variants in *RAD51D* and *XRCC2*, which encode RAD51 paralogs that form a heterodimer within the BCDX2 complex, confer increased cancer risk and homologous recombination deficiency. However, most *RAD51D* and *XRCC2* variants in ClinVar are classified as variants of uncertain significance (VUS), limiting clinical utility. Here, we applied saturation genome editing (SGE) to measure the effects of 5,412 *RAD51D* and 3,743 *XRCC2* variants on cellular fitness and additionally assessed 2,876 and 2,069 variants in these genes for effects on RNA expression. Fitness scores discriminated pathogenic from benign variants with near-perfect accuracy (AUC=0.994 for *RAD51D*; AUC=1.000 for *XRCC2*). Integration of RNA expression data revealed *RAD51D*, but not *XRCC2*, is exceptionally sensitive to splice-altering variation, with 24% of RAD51D loss-of-function missense variants acting through RNA-mediated mechanisms compared to only 5% in *XRCC2*. These SGE datasets provide strong, splice-resolved functional evidence to support variant classification across both genes.

## Introduction

Germline pathogenic variants in *RAD51D* confer significantly increased risk of ovarian cancer (OR∼7)^1,2,3^ and breast cancer^4,5^, with a particularly strong association with triple-negative disease (OR=6.97)^6^. *RAD51D* mutation carriers are more likely to develop estrogen receptor (ER)-negative, high-grade tumors compared to non-carriers^7,8^. While the cancer risk associated with monoallelic *XRCC2* variants remains less well-defined^9,10,11^, biallelic pathogenic variants in this gene cause Fanconi anemia, a severe disorder characterized by bone marrow failure and cancer predisposition^12,13^. Given their critical roles in DNA repair and cancer risk, *RAD51D* and *XRCC2* are included on hereditary cancer testing panels^14,15,16^. However, of the 1,545 single nucleotide variants (SNVs) in *RAD51D* and 601 SNVs in *XRCC2* reported in the January 2025 ClinVar release, >50% and >60% respectively are classified as variants of uncertain significance (VUS). The vast majority of missense variants are VUS, with 82% (755 of 920) and 93% (375 of 405) for *RAD51D* and *XRCC2* respectively. This high VUS burden limits accurate genetic counseling and risk assessment. Furthermore, loss of *RAD51D* results in homologous recombination deficiency (HRD), rendering *RAD51D*-deficient tumors sensitive to PARP inhibitors and platinum-based chemotherapies^1,17,18,19^. While RAD51D-XRCC2 function together, clinical evidence for *XRCC2* deficiency and HRD has been reported but remains limited^20,21,22^. Comprehensive variant-level functional data is essential both for cancer risk assessment and for identifying patients whose tumors are most likely to respond to targeted therapies.

*RAD51D* and *XRCC2* encode RAD51 paralogs that form an obligate heterodimer and act with other RAD51 paralogs to regulate RAD51 filament formation. Together, the RAD51 paralogs form the BCDX2 complex (RAD51B-RAD51C-RAD51D-XRCC2)^23,24^ and a recently described XRCC3 complex (XRCC3-RAD51C-RAD51D-XRCC2)^25,26^. During homologous recombination, BCDX2 binds RPA-coated ssDNA and facilitates RPA-to-RAD51 exchange, while the XRCC3 complex caps and stabilizes the 5’ end of RAD51 filaments. Together, these complexes enable RAD51 presynaptic filament formation required for homology search and strand exchange^27,24,23,28^. Consistent with their essential roles in this pathway, loss of RAD51D or XRCC2 renders cells hypersensitive to DNA-damaging agents and abolishes RAD51 focus formation following DNA damage^29^. While studies have begun to identify specific residues essential for heterodimer formation and HR activity^30,24,23^, comprehensive variant-level functional mapping across the full RAD51D-XRCC2 interface in a clinical classification context has only recently been undertaken^31^.

Saturation genome editing (SGE) enables systematic functional assessment of thousands of variants in genes essential for the growth of haploid HAP1 cell line^32,33^. By introducing all possible SNVs and 3 base-pair (bp) deletions into the endogenous locus of haploid cells and tracking variant abundance over time, SGE measures variant effects on both cell fitness and RNA expression. This approach has been successfully applied to multiple genes in the HR pathway, including *BRCA1*^33,34^, *BRCA2*^35,36^, *BARD1*^37^, *BAP1*^38^, *PALB2*^39^, and *RAD51C*^40^, generating comprehensive functional maps that accurately discriminate pathogenic from benign variants. Like other HR pathway members, *RAD51D* and *XRCC2* are essential for HAP1 cell survival, making them amenable to SGE analysis.

Here we measured the effect of 5,412 *RAD51D* variants and 3,743 *XRCC2* variants on cellular fitness using SGE, and additionally measured the effect of 2,876 *RAD51D* and 2,069 *XRCC2* variants on RNA expression. Fitness scores discriminated pathogenic from benign variants with near-perfect accuracy for both genes (AUC=0.994 for *RAD51D*; AUC=1.000 for *XRCC2*), and correctly identified all known pathogenic missense variants as loss-of-function (LoF). Integration of RNA abundance measurements revealed that *RAD51D* is exceptionally sensitive to splice-altering variation with 24% of LoF missense variants acting through RNA-mediated mechanisms, a rate 2-6 fold higher than reported for other genes^33,34,37,41^. We find that the transcript architecture of *RAD51D* likely underlies this sensitivity. Furthermore, we demonstrate that the protein sequence encoded by the C-terminal exon of *RAD51D* is dispensable for function, with implications for the automatic classification of truncating and canonical splice variants in this region as pathogenic. Mapping LoF missense variants onto solved structures revealed that the RAD51D-XRCC2 heterodimer interface is broadly tolerant to missense variation caused by single nucleotide variants and that apparent interface residues such as RAD51D p.Lys297 are critical for ATP coordination rather than protein-protein interaction. Combined with computational variant effect predictions, SGE provides sufficient evidence to reclassify 87% *RAD51D* VUS and 98% *XRCC2* VUS, demonstrating immediate utility for hereditary cancer risk assessment and treatment decisions.

## Results

### Fitness scores for 5,412 *RAD51D* and 3,743 *XRCC2* variants generated by saturation genome editing

To perform SGE on *RAD51D* and *XRCC2* we designed sgRNAs and libraries of repair templates to edit all possible SNVs and 3-base pair (bp) deletions across all 10 coding exons and proximal flanking introns of *RAD51D* (4,524 designed SNVs and 1,131 3-bp deletions) and the three coding exons of *XRCC2* (3,153 designed SNVs and 788 3-bp deletions) (**Fig. 1 a,b**). We validated sgRNAs and generated 14 repair template libraries for *RAD51D* and 11 for *XRCC2*, with each library covering a single exon or sub-exon region (**Fig. 1b**; **Extended Data Fig. 1a,d**). SGE experiments were performed in triplicate in HAP1-Δ*LIG4* cells and sampled at days 5, 13 and 17 for *RAD51D* and days 5 and 13 for *XRCC2* (**Fig. 1c**). Genomic DNA was sequenced at each timepoint to an average depth of >2,000 reads per variant. Editing rates for generating usable sequences containing a single variant and programmed Cas9-blocking edits ranged from 11.5% to 39.8% for *RAD51D* and from 11.1% to 32.4% for *XRCC2* (**Extended Data Fig. 1b,e**). Variant counts were highly reproducible across experimental replicates (*RAD51D* median Pearson *r*=0.89, range 0.61-0.99; *XRCC2* median Pearson *r*=0.90, range 0.53-0.98) (**Extended Data Fig. 1c,f**).

**Figure 1.**
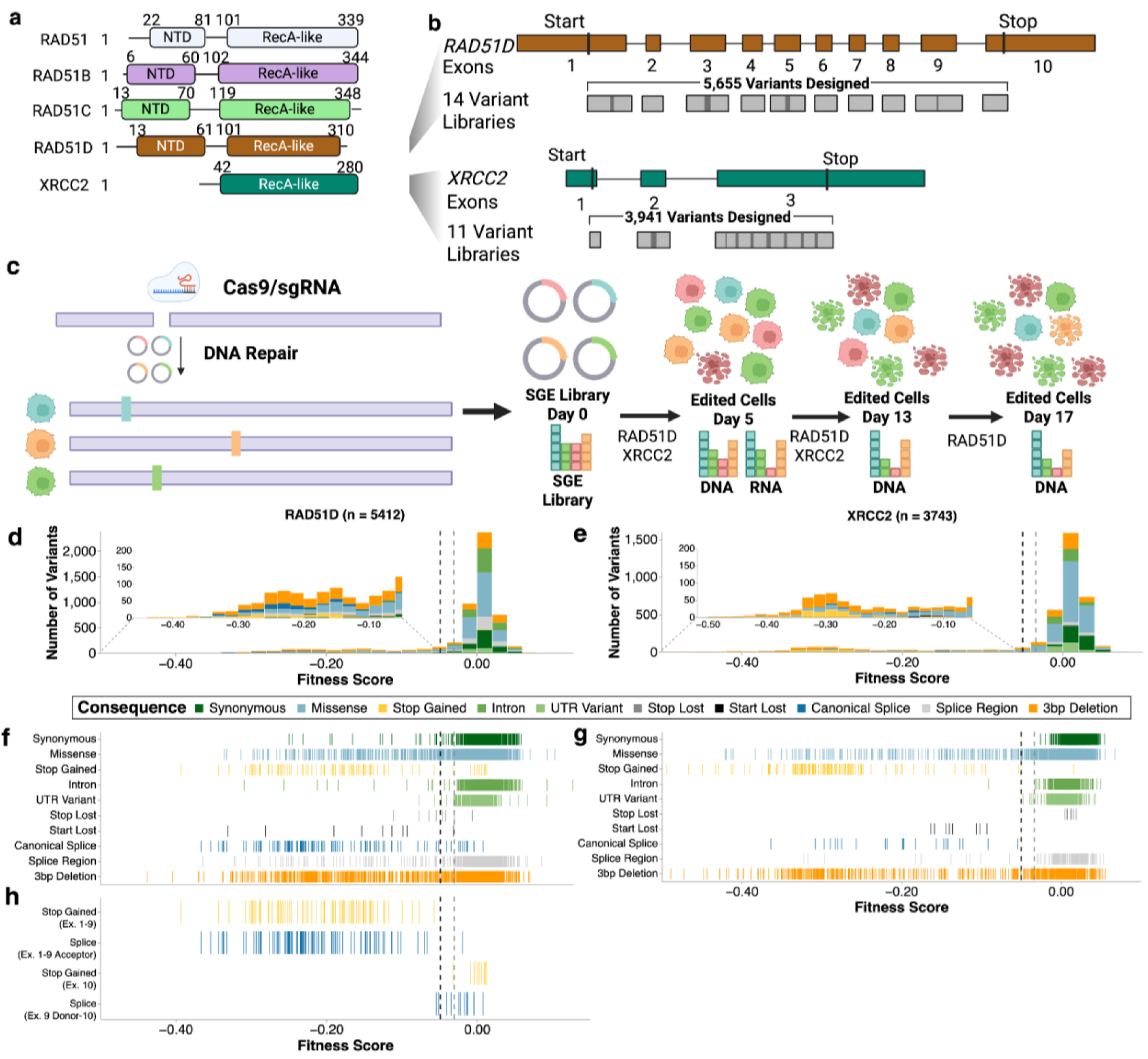
Saturation Genome Editing to functionally assess 9,155 variants of *RAD51D* and *XRCC2*. a) RAD51 (light blue), RAD51B (purple), RAD51C (green), RAD51D (brown), and XRCC2 (teal) domain structure. b) Maps of exon structure followed by coverage of repair template libraries required to saturate all coding exons and proximal introns for *RAD51D* (top) and *XRCC2* (bottom). 14 variant libraries were required for *RAD51D* and 11 for *XRCC2*. Exons that were too large for a single repair template library required sub-exon-sized repair template libraries and paired sgRNAs. c) Schematic of SGE experimental strategy and scoring process. For *XRCC2*, two timepoints (day 5 and 13) were collected while for *RAD51D*, three time points (days 5, 13, and 17) were collected. At all timepoints, DNA was sampled. On day 5, for *RAD51D*, RNA was also sampled. d) Histogram of fitness scores for 5,412 *RAD51D* variants colored by mutational consequence as indicated. Vertical black and grey lines at X=-0.0487 and X=-0.0304 represent thresholds for the functionally abnormal and functionally normal classes respectively. Inset highlights functionally abnormal variants. e) Histogram of fitness scores (n=3,743) for *XRCC2*. Vertical black and grey lines at X=-0.0498 and X=-0.0335 represent thresholds for the functionally abnormal and functionally normal classes respectively. f) Strip plot of *RAD51D* fitness scores organized by mutational consequence (n=5,412). g) Strip plot of *XRCC2* fitness scores organized by mutational consequence (n=3,743). h) Strip plot of *RAD51D* fitness scores highlights canonical splice and stop-gained variants (n=261) at the exon 9 and 10 splice junction and in exon 10 respectively.

Fitness scores were calculated by modeling variant depletion over time with a linear model^37^ (see methods and IGVF Assay Standards). We generated fitness scores for 4,340 SNVs (>95% of designed) and 1,072 3-bp deletions (>90% of designed) for *RAD51D*, and 3,023 SNVs (>95% of designed) and 720 3-bp deletions (>90% of designed) for *XRCC2* (**Fig. 1d,e; Extended Data Fig. 2a,b; Supplementary File 1, 2**). All data is available on MaveDB and the IGVF data portal. For both genes, fitness scores for synonymous and intronic variants were largely centered around zero, representing no change in abundance over the time course (median score: *RAD51D*=0.005, *XRCC2*=0.007), while stop-gained and canonical splice variants showed strong depletion (**Fig. 1f,g**). We determined thresholds to classify variants as LoF or functionally normal by modeling the distributions of variants expected to be damaging (stop-gained) and variants expected to be tolerated (synonymous and intronic) using a Gaussian Mixture Model (**Fig. 1d,e** dotted lines; **Extended Data Fig. 3**).

For *RAD51D*, 82/95 (86.3%) of stop-gained variants and 149/166 (89.8%) of canonical splice variants were classified as LoF. All 13 stop-gained variants in the terminal coding exon (exon 10) scored as functionally normal or indeterminate and 15 of 18 canonical splice site variants within the exon 9-10 junction were functionally normal or indeterminate (**Fig. 1h**), suggesting the coding sequence of exon 10 is dispensable for RAD51D function. As expected, most synonymous variants (612/637, 96.1%) and intronic variants (984/1,001, 98.3%) were functionally normal. Among missense variants, 227/1,874 (12.1%) were classified as LoF, and 96/686 (14.0%) of splice region variants were LoF.

For *XRCC2*, 143/144 (99.3%) of stop-gained and 36/37 (97.3%) of canonical splice variants were LoF. 473/473 (100%) of synonymous and 385/386 (99.7%) of intronic variants were functionally normal. Among missense variants, 172/1,693 (10.2%) were classified as LoF, a proportion similar to *RAD51D* (*p*=0.070, Fisher’s exact test) consistent with their structural homology (**Fig. 1f,g**). However only 11/159 (6.9%) splice region variants were LoF which is significantly lower than *RAD51D* (14.0%, p=0.017, Fisher’s exact test) suggesting *RAD51D* may have increased sensitivity to splice-altering variation compared to XRCC2.

### Deletion maps and LOF missense variants identify residues of RAD51D and XRCC2 required for function

Mapping the fitness scores of 3-bp deletions to the RAD51D protein sequence shows that deletions that result in stop-gained codons are LoF only in the first 300 amino acids (**Fig. 2a**), reinforcing our results showing that amino acids 301-329 encoded by exon 10 are dispensable for function (**Fig. 1h**). LoF 3-bp deletions that do not result in stop-gained codons are spread throughout amino acid positions 1-300 including in the N-terminal and RecA domains suggesting amino acids in these domains are required for function. This is mirrored by LoF missense variants which are also found throughout positions 1-300 (**Fig. 2a**) and enriched at the Walker A (aa 107-114) and B (aa 198-206) motifs (2.06-fold; p=1.92e-04, Fisher’s exact test). The sole LoF missense variant in exon 10, p.Thr303Lys; Thr303 contacts AMP-PNP at the XRCC2–RAD51D interface of the secondary ATP binding site^23^.

**Figure 2.**
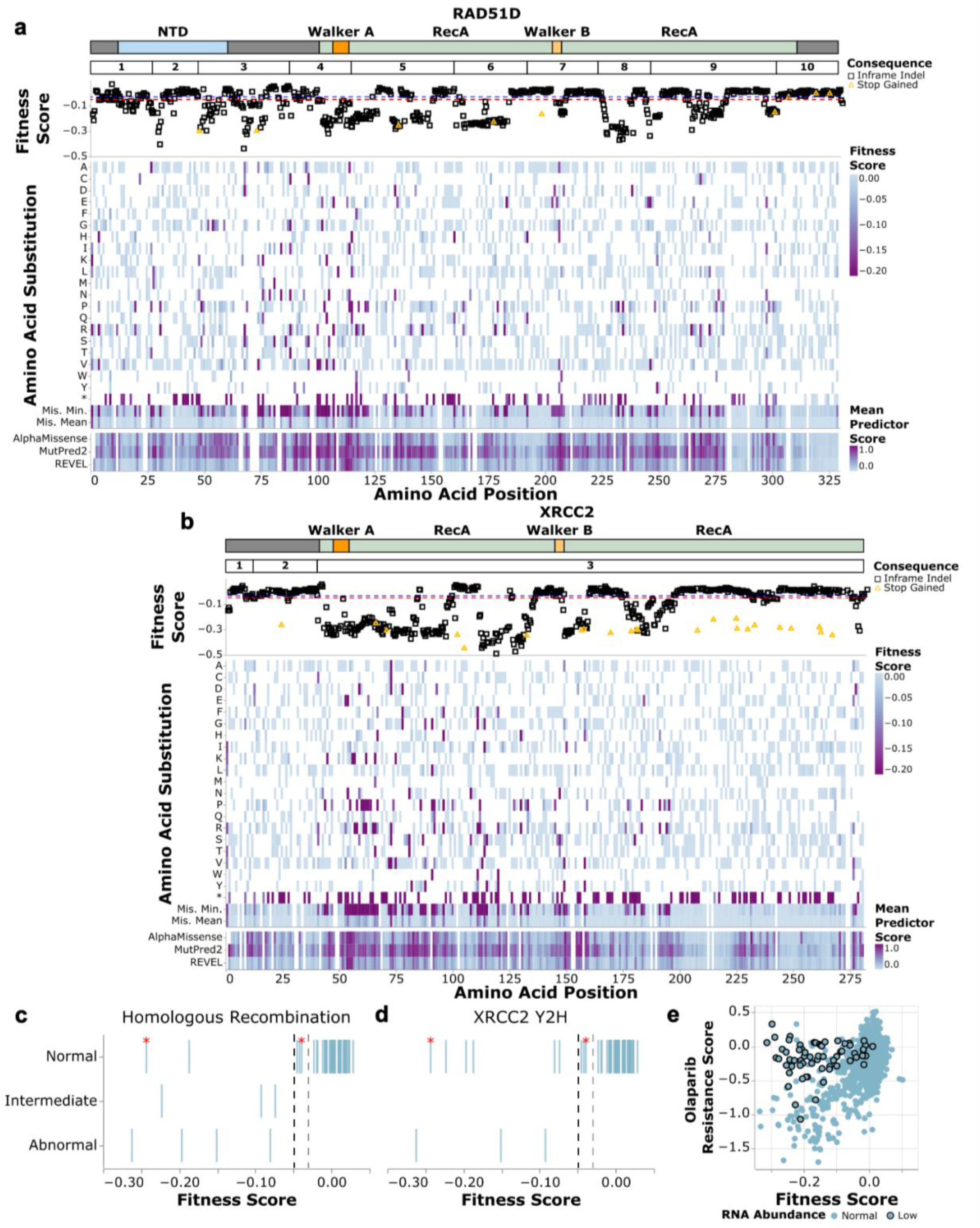
SGE reveals protein residues required for function. a) Map of programmed 3-bp deletions in *RAD51D* (n=1,072) (top). Amino acid positions in RAD51D are on the X-axis and fitness scores are on the Y-axis. In-frame deletions and deletions that cause stop-gained variants are denoted by shape and color. A heatmap of fitness scores for missense and stop-gained variants (middle). Amino acid positions in RAD51D are on the X-axis and amino acid and stop-gained (*) substitutions are indicated on the Y-axis. Fitness scores are colored as indicated. Heatmap of minimum and mean fitness scores across missense variants and mean variant effect predictor scores (bottom). For predictors, high scores denote predicted pathogenicity, while low fitness scores indicate LoF. b) Analogous figure as in (a) for *XRRC2* 3-bp deletions (n=720),missense and stop-gained variants. c) and d) Strip plots of *RAD51D* variants assessed in orthogonal assays for homologous recombination (c; n=62) and yeast 2-hybrid (Y2H) interaction with *XRCC2* (d; n =62)^31^. Fitness score is on the X-axis and functional consequence of the variant in its respective biochemical assay is on the Y-axis. Variants with decreased RNA (RNA score <-2.79) are highlighted with a red asterisk. d) Olaparib resistance score^31^ (Y-axis) vs SGE fitness score (X-axis) for missense variants assayed in both studies (n=1,871). Variants with low RNA are outlined in black.

In XRCC2, 3-bp deletions that result in stop-gained codons are LoF and are found throughout the full length of the protein suggesting that the protein cannot be truncated and retain function (**Fig. 2b**). However, in-frame deletions and missense variants are generally tolerated at amino acid positions 2-40 encoded by exons 1 and 2 N-terminal to the RecA domain and between amino acids 200-275 suggesting no single amino acid in those regions is required for function. Most LoF missense variants occur between residues 41 and 195 in the RecA domain between the Walker A and Walker B motifs (p=6.538 e-26, Fisher’s exact test).

Comparison of fitness scores to variant effect predictors AlphaMissense^42^, MutPred2^43^, and REVEL^44^ shows general concordance with functional classifications (**Fig. 2a,b**; **Extended Data Fig 4a-j**). Across both genes, the predominant form of discordance was overprediction of pathogenicity (**Extended Data Fig. 4a-d, f-i**). For the majority of stop-gained variants, CADD scores^45,46^ and SGE functional classifications were concordant. However, in *RAD51D*, 7 of 86 stop-gained variants (8.1%) had high CADD deleteriousness scores (>30) yet were functionally normal by SGE, all occurring in exon 10 (**Extended Data Fig. 4e**). Conversely, in XRCC2, 8 of 121 stop-gained variants (6.6%) had abnormal SGE scores despite CADD scores <30 (**Extended Data Fig. 4j**). This highlights the limitations of computational predictors as standalone tools and underscores the value of experimentally derived functional measurements.

To validate that fitness scores measured by SGE integrate known cellular functions of RAD51D and XRCC2, we compared them to results from orthogonal functional assays (**Fig. 2c-d**). Orthogonal functional assays are scarce for XRCC2 variants, but three of the four missense variants found to have modest defects in HR activity in human cells were also LoF by SGE and all variants with normal HR activity were functionally normal^47^ (**Extended Data Fig. 5a**). Fitness scores for RAD51D were highly concordant with orthogonal measurements of HR activity (54/62 missense variants; 87.1%) and XRCC2 interaction (53/62; 85.5%) (**Fig. 2c,d**). We also compared our SGE fitness scores to an independent MAVE dataset for RAD51D^31^. In this MAVE, a U2OS cell line engineered to lack endogenous *RAD51D* was used to assess a library of amino acid substitutions expressed from cDNA for resistance to PARP inhibition by olaparib. Despite differences in cell type, assay readout, and experimental design, these two multiplexed assays were remarkably concordant across all variant types (n=2,456; r=0.599, p=9.28e-239) and specifically for missense variants (n=1,871; r=0.515, p=5.053e-127) (**Fig. 2e; Extended Data Fig. 5b**). For each of these three orthogonal assays there were variants where the endogenously expressed variants assessed by SGE were LoF while the variants expressed from cDNAs in the orthogonal assay were functionally normal. Mapping our RNA scores to these variants, shows that 61% (30 of 49) missense variants that were LoF in the SGE but not in the cDNA-based assays had low RNA scores suggesting these variants cause splice defects that cDNA-based assays cannot measure (**Fig. 2e; Extended Data Fig. 5b**). Thus, SGE-derived fitness scores for RAD51D and XRCC2 integrate the functions of both proteins in HR and RAD51D’s interaction with XRCC2 while additionally capturing variants that impact RNA expression.

### Integrating variants effects on RNA expression and cellular fitness reveals extreme sensitivity of *RAD51D* to splice-altering variation

To distinguish variants that disrupt protein function from those that affect RNA expression or splicing, we used targeted RNA sequencing to quantify transcript levels following SGE of *RAD51D* and *XRCC2* (**Fig. 1c**). For each SGE experiment, RNA was collected 5 days post-transfection for each replicate, reverse-transcribed using a gene-specific primer in the 3’UTR and amplified with intron-spanning primers to exclude genomic DNA background. Amplicons were aligned to the MANE Select transcripts (*RAD51D*: NM_002878.4; *XRCC2*: NM_005431.2) to assign variants and RNA variant counts across replicate experiments were highly concordant (*RAD51D* RNA median Pearson *r*=0.93, range 0.80-0.98; *XRCC2* RNA median Pearson *r*=0.86, range 0.70-0.92; **Extended Data Fig. 6a,c**). RNA scores were calculated as the log_2_ ratio of variant RNA abundance to genomic DNA abundance.

We generated RNA scores for 2,876 *RAD51D* variants comprising 2,655 coding and 221 untranslated region variants (**Fig. 3a**) and for 2,069 *XRCC2* coding region variants (**Fig. 3b**). For *RAD51D*, RNA and fitness scores for all variants showed a moderate positive correlation (Pearson’s *r*=0.502, p=2.279e-183, 95% CI [0.474, 0.529) (**Fig. 3c, Extended Data Fig. 6b,d**), with variants in exons 3 and 4 showing substantially stronger correlations (*r=*0.819, p=2.377e-70, 95% CI [0.777, 0.854] and *r*=0.653, p=5.277e-28, 95%CI [0.570, 0.723], respectively) indicating that transcript abundance accounts for 67.1% and 42.6% fitness score variance in these exons. Using a threshold for low RNA abundance at 3 standard deviations below the mean (-2.79 log_2_ fold-change), 86% (80/93) of variants below this threshold were LoF (**Extended Data Fig. 6b,d**). Of the 13 variants retaining normal or indeterminate fitness scores despite having low RNA abundance, 7 were located at or near the exon 9-exon 10 junction, consistent with functional dispensability of *RAD51D* C-terminus (**Fig. 3a, Fig.1h**) and of the remaining 6, four had SpliceAI predictions^48^ consistent with altered splicing resulting in in-frame deletions. Overall, 23.2% (80/345) of all LoF *RAD51D* variants had low RNA scores including 54 of the 227 (23.8%) LoF missense variants, while 99.6% (2,450/2,460) of functionally normal variants had normal RNA scores (**Fig. 3c**). For *XRCC2*, RNA and fitness scores showed a weak negative correlation (Pearson’s r=-0.348, p=4.991e-60, 95% CI [-0.385,-0.310]), likely reflecting both a low sensitivity to altered splicing as well as the overall absence of NMD (**Extended Data Fig. 6d**).

**Figure 3.**
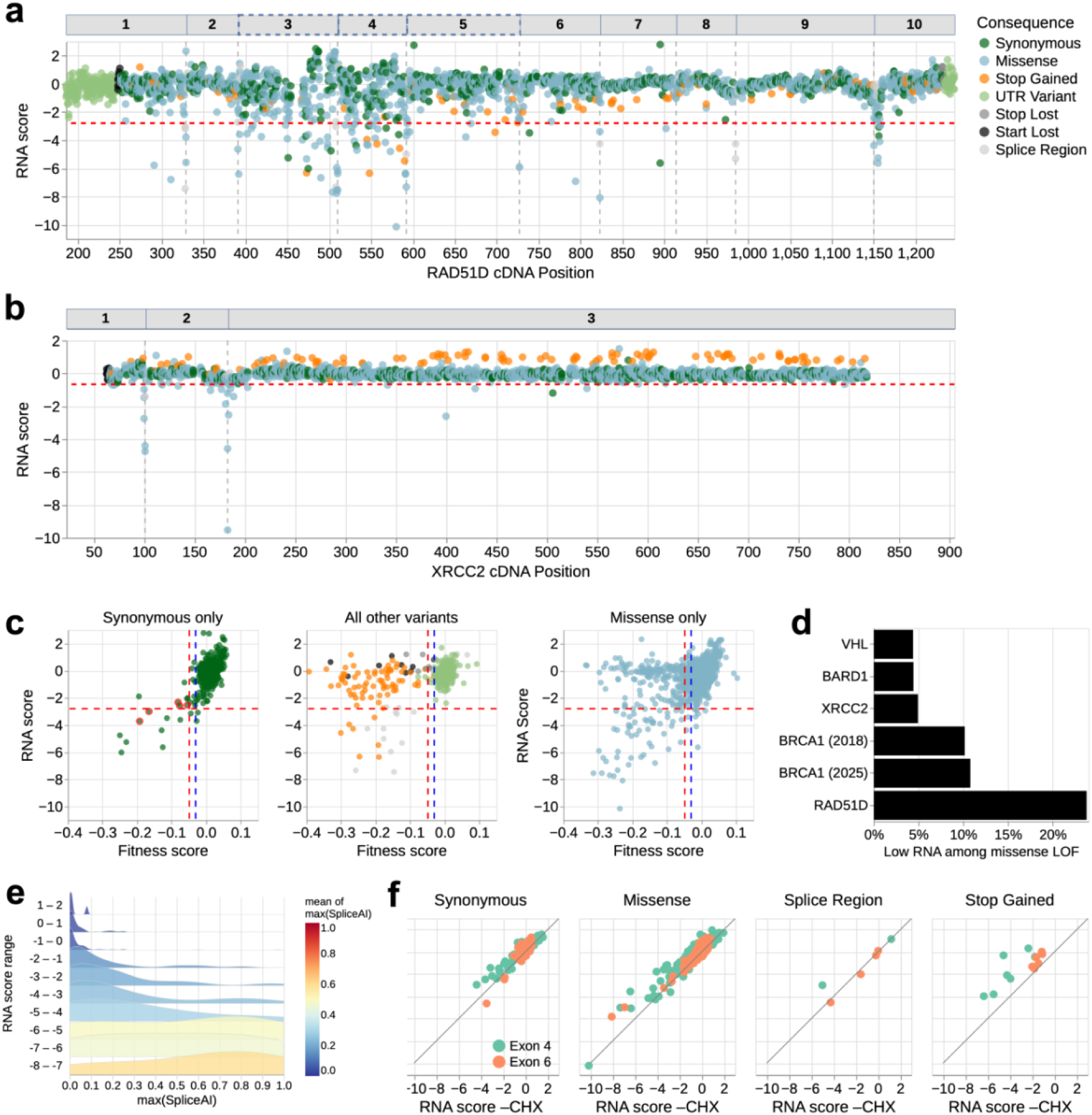
RNA scores identify LoF variants acting through RNA-level effects. (a) RNA scores for all *RAD51D* variants (n=2,876) plotted by cDNA position. The transcript exon structure is depicted above the plot with exon boundaries indicated by vertical dashed lines. Variants are colored by molecular consequence. The horizontal red dashed line denotes the threshold for low RNA (-2.79 log2 fold-change). (b) RNA scores for all *XRCC2* variants (n=2,069) plotted by cDNA position, analogous to (a). The horizontal red dashed line denotes the threshold for low RNA abundance (-0.66 log2 fold-change). (c) Scatterplot of *RAD51D* fitness scores (X-axis) versus RNA scores (Y-axis) for *RAD51D* synonymous variants (left), for all other variants (center), and for missense variants only (right) colored by molecular consequence. Five synonymous variants with LoF fitness scores but likely benign in ClinVar have low or near-threshold RNA scores (c.225G>A, c.243C>T, c.300A>G, c.321A>G, c.333C>T; red outline). Three of these variants (c.300A>G, c.321A>G, c.333C>T) also have high SpliceAI, providing orthogonal computational support for splice disruption. The red dashed horizontal line indicates the low RNA score threshold, and the blue and red dashed vertical lines indicate SGE thresholds for normal and LoF fitness scores. (d) Bar chart showing the proportion of LoF missense variants with low RNA scores in *RAD51D* compared to *XRCC2* and four published SGE datasets (*BARD1*^37^, *BRCA1*^33,34^, and *VHL*^41^). (e) Ridgeline plot of SpliceAI maximum scores (X-axis) across bins of RNA abundance scores (Y-axis) for functionally abnormal *RAD51D* variants only. Colors indicate the mean SpliceAI maximum score per bin ranging from low (blue) to high (red). (f) Comparison of RNA scores generated with and without cycloheximide (CHX) treatment for synonymous, missense, splice region, and stop-gained variants for *RAD51D* exons 4 and 6.

To establish whether the high proportion of missense variants acting through RNA-level mechanisms is a gene-specific property of *RAD51D*, we compared LoF missense variants with low RNA across four SGE datasets in which RNA was measured alongside cellular fitness. The low RNA scores for the 54 *RAD51D* LoF missense variants suggests that these variants impair *RAD51D* function through RNA-mediated rather than protein-mediated effects. This proportion of LoF missense variants was approximately two-fold higher than for *BRCA1* (10%-11%^33,34^) and approximately five-to six-fold higher than for *XRCC2* (5%), *BARD1*^37^(4%) and *VHL*^41^ (4%) (**Fig. 3d**). These results suggest that *RAD51D* susceptibility to RNA-mediated loss of function is gene-specific and underscore the value of integrating RNA abundance measurements into SGE experimental designs. Of the 54 RNA-mediated LoF missense variants, only 29 (53.7%) had SpliceAI scores ≥0.20 suggesting splice site disruptions as the underlying mechanism; the remaining 25 had low SpliceAI predictions and were enriched in exons 3 and 4 (20/54), suggesting these exons may be particularly sensitive to splice disruption through mechanisms not captured by state-of-the-art splice prediction tools^48^ (**Extended Data Fig. 6a**).

To determine whether the enrichment of RNA-mediated LoF variants in exon 3 and 4 could be explained by the transcript structure of *RAD51D*, we examined its isoform expression profile of our HAP1-Δ*LIG4* cell line. In our line, the full length transcript constituted 27% of total *RAD51D* expression as measured by RT-PCR and long read RNA sequencing, comparable to the mean proportions among GTEx breast and ovary samples (25.9% and 26.0%, respectively)^49,50^ **(Extended Data Fig. 6e,f**). The predominant alternative isoforms lack exons 3, 4, and 5 or exon 3, and both are reported to be loss-of-function for HR^51,52,53,30^. This transcript architecture may explain the observed sensitivity: because the full-length transcript is already a minority of total *RAD51D* expression in both HAP1 cells and disease-relevant tissues, any splice-disrupting variant that further reduces its abundance may be sufficient to cause loss-of-function. Of 248 ClinVar variants in exons 3 and 4, 164 are classified as VUS (97 in exon 3, 67 in exon 4). Among these, 17 VUS have low RNA scores, of which 15 are missense variants (8 in exon 3, 7 in exon 4) impairing function through RNA-mediated splice disruption, underscoring the clinical relevance of this transcript architecture.

RNA scores provided mechanistic insight across all variant types and revealed the limitations of SpliceAI as a standalone predictor. In *RAD51D*, only 55% (44/80) of LoF variants with low RNA had SpliceAI ≥0.20, and among 16 LoF synonymous variants (2.5% of 637 total, predominantly in exons 3 and 4), SpliceAI predicted altered splicing in only 4 (25%) whereas 7 (44%) had low RNA scores without evidence of novel splice site creation, possibly implicating alternative splicing mechanisms, such as exonic splicing regulatory elements, beyond those captured by computational prediction (**Fig. 3c**). A consistent pattern was observed in XRCC2, where 78% (7/9) of LoF missense variants with low RNA had SpliceAI ≥0.20 (**Extended Data Fig. 6g**). However, across both genes, only 4% of functionally normal *RAD51D* variants (n=2,450) and 2.8% of functionally normal *XRCC2* variants (n=1,745) had SpliceAI scores ≥0.20 reflecting a low overall false-positive rate (**Fig. 3e, Extended Data Fig. 6g)**.

### RNA scores reveal regional sensitivity to nonsense-mediated decay for *RAD51D* and unexpected start-and stop-loss effects

Stop-gained variants in *RAD51D* showed unexpectedly high rates of escape from nonsense-mediated decay (NMD): only 7 of 54 (13.0%) variants expected to be targeted by NMD triggered transcript degradation. Stop-gained variants that triggered NMD were significantly enriched in exons 3–5 (28% vs 0%; p=2.714e-03, Fisher’s exact test) (**Fig. 3a,c; Extended Data Fig. 6h**). To confirm that reduced RNA abundance reflected NMD for stop-gained variants, we treated cells with cycloheximide (CHX) after performing SGE on exons 4 and 6 to inhibit NMD and compared RNA scores across variant types (**Fig. 3f)**. Stop-gained variants in both exons shifted toward higher RNA scores upon CHX treatment, but exon 4 variants showed significantly greater CHX rescue than those in exon 6 (∼2.8 fold, p=0.0003, Mann-Whitney U test). These results are consistent with regional NMD sensitivity and suggest that stop-gained variants in exons unique to the full-length transcript are more susceptible to NMD than stop-gained variants elsewhere in the gene. Compared to other SGE datasets with RNA scores, and with the exception of *XRCC2* that has no NMD eligible variants and *VHL* where all NMD eligible variants escape NMD, *RAD51D* has the highest proportion of NMD-escaping stop-gained variants across all genes examined (47/54, 87.0%; **Extended Data Fig. 6h**).

start-and stop-lost variants reveal distinct mechanisms by which translation can be affected independent of splicing or NMD. While all start-loss variants of XRCC2 were LoF, 2 of 14 RAD51D start codon (Met1) variants did not cause loss-of-function, consistent with the strong Kozak consensus context (positions −3, −1, +4: A, C, G) and the absence of viable downstream alternate start codons; Met16, the nearest downstream methionine, has a weak Kozak context lacking the critical −1 and +4 requirements^54,55^. The two functionally normal variants illustrate distinct compensatory mechanisms: c.-2_1delACA (p.Met1=) maintains methionine while disrupting the Kozak sequence at position −1, and c.2T>C (p.Met1Thr) creates a non-canonical ACG start codon that supports translation via Met-tRNAi, incorporating methionine rather than threonine^56,57^ (**Extended Data Fig. 7a**). RAD51D c.1A>T (p.Met1Leu), reported in individuals affected with breast and ovarian cancer ^58,59^, was recently identified in a small Danish family in which the proband’s mother carried the variant and developed ovarian cancer at age 50 (**Extended Data Fig. 7b**).

Stop-loss variants create C-terminal peptide extensions by translational read-through into the 3’ UTR. All *XRCC2* stop-loss variants were functionally normal, but 10 of 12 *RAD51D* stop-loss variants scored as LoF or intermediate **(Fig. 1f**). The *RAD51D* stop codon (Ter329) is followed by 168 bp of 3’ UTR sequence before encountering another downstream stop codon, predicting a 56-amino-acid extension and avoiding non-stop decay^60^. LoF stop-loss variants were an unexpected finding given the dispensability of the C-terminus of RAD51D, however the extended tail is predominantly hydrophobic (56% hydrophobic residues, including 11 leucines), with hydrophobicity increasing toward the C-terminus (73% in the final 15 residues) suggesting this extension could trigger protein degradation^61,62^ **(Extended Data Fig. 7c**).

### LoF missense variants reveal structural insights into RAD51D and XRCC2 domains

While some LoF missense variants reduced RNA abundance, others likely disrupted protein function or stability; we therefore mapped fitness scores onto solved structures to explore sequence-function relationships.

Residues in the critical Walker A and B ATP-binding motifs in RAD51D and XRCC2 were generally intolerant to missense changes, particularly those with major physicochemical changes (**Fig. 2a-b**; **Fig. 4a-b, 4e; Extended Data Fig. 8a; Extended Data Fig. 9a-b**). Beyond the Walker A and B motifs, RAD51D p.Arg145 and XRCC2 p.Arg91 may form non-bonded interactions with ATP^23–26^. However, RAD51D p.Arg145 was generally tolerant to missense variants while XRCC2 p.Arg91 was wholly intolerant, suggesting differing roles in ATP interaction (**Fig. 4e, Extended Data Fig. 8a**).

**Figure 4.**
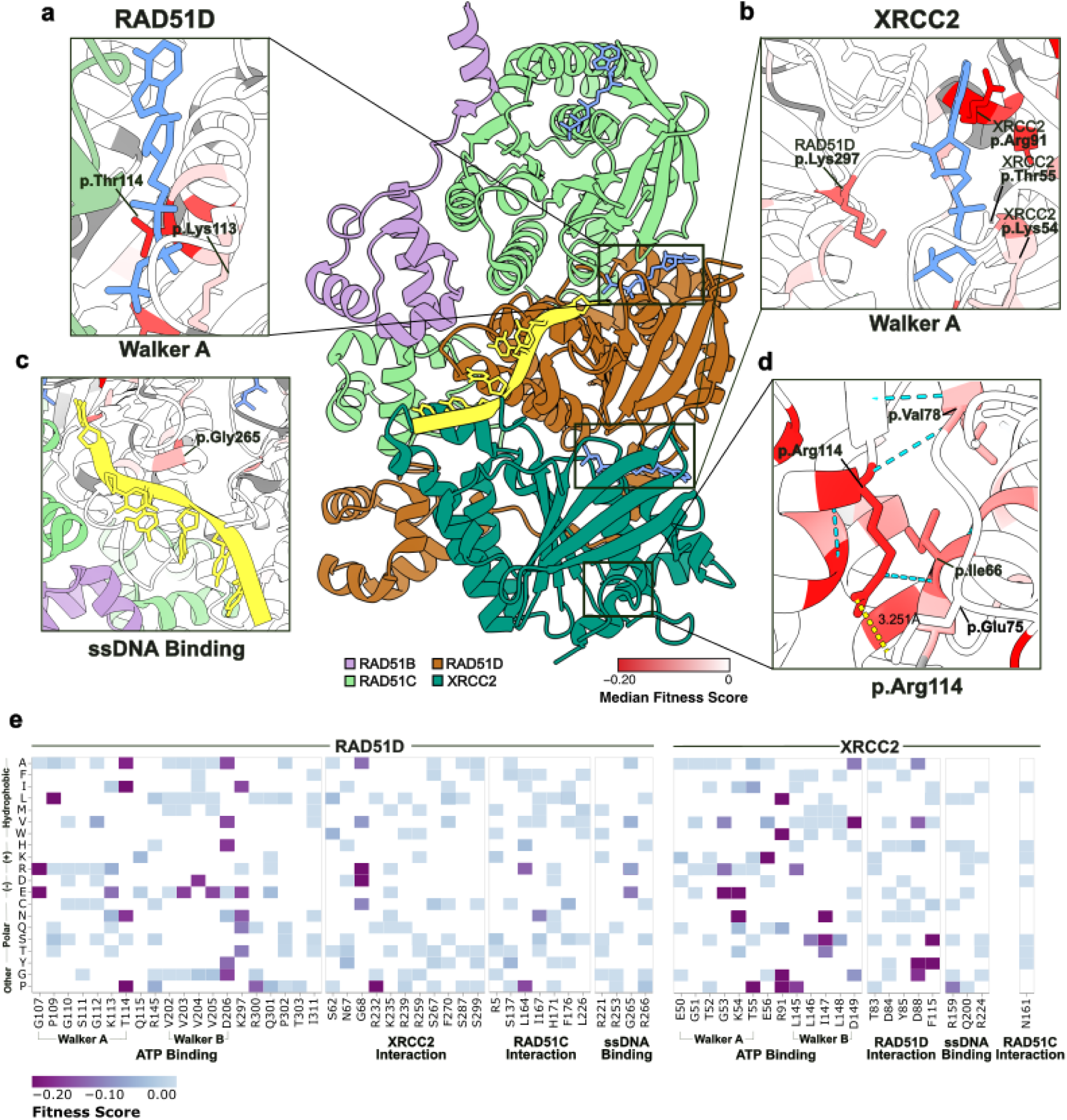
Functional and structural consequences missense variants on RAD51D and XRCC2. (a) Diagram of RAD51D’s Walker A ATP-binding motif (PDB: 8GBJ^23^) colored by median fitness score as indicated (gray denotes residues not scored). RAD51D p.Lys113 and p.Thr114 are highlighted. RAD51C is colored in green and ATP is colored in blue. (b) Diagram of XRCC2’s Walker A ATP-binding motif (PDB: 8GBJ^23^) colored by median fitness score. RAD51D p.Lys297 and XRCC2 p.Lys54, p.Thr55, and p.Arg91 are highlighted. ATP is colored in blue. (c) ssDNA binding motif of RAD51D (PDB: 8GBJ^23^) colored by median fitness score. Bound ssDNA is colored in yellow. RAD51B and RAD51C are colored in purple and green respectively. RAD51D p.Gly265 is highlighted. (d) XRCC2 p.Arg114 (PDB: 8GBJ^23^) and potential interacting partners colored by median fitness score. Dotted yellow line denotes an electrostatic interaction and dotted blue lines highlight potential hydrogen bonding interactions. (e) Heatmap of residues previously reported^23,24^ in RAD51D (left) and XRCC2 (right) biochemical interactions. Y-axis denotes substituted amino acids. X-axis denotes original amino acid and residue position. Residues are grouped by their biochemical function. Fitness scores are colored as indicated.

During DNA repair, RAD51D and XRCC2 interact with ssDNA^23–26,63^, but residues implicated in this interaction were generally tolerant to missense variation, except for RAD51D p.Gly265 and XRCC2 p.Arg159 (**Fig. 4c, 4e, Extended Data Fig. 8a, Extended Data Fig. 9c**). Only p.Gly265Ala was tolerated in RAD51D, suggesting a small hydrophobic residue is required for the local structure supporting ssDNA binding (**Fig. 4c**), while p.Arg159Pro was not tolerated in XRCC2, likely disrupting helical secondary structure (**Fig. 4e**). Notably, arginine residues potentially forming charged interactions with ssDNA were broadly tolerant to charge-disrupting substitutions (**Fig. 4e, Extended Data Fig. 8a-b**), and residues near ssDNA in RAD51D (p.263–267) similarly tolerated alanine substitutions in an Olaparib resistance assay^31^. Together, these findings indicate that successful ssDNA binding depends on correct local structure and multi-residue interactions rather than any single critical residue.

At the RAD51D–XRCC2 and RAD51C interfaces, most RAD51D–XRCC2 interface residues were missense tolerant, except RAD51D p.Lys297 and XRCC2 p.Asp88 (**Fig. 4e, Extended Data Fig. 8a, 8c**). As p.Lys297 sits at a shared interface with ATP, and only the charge-preserving p.Lys297Arg was tolerated - consistent with the Olaparib resistance assay^31^ - it likely plays a greater role in ATP coordination than XRCC2 binding. This is further supported by the missense tolerance of XRCC2 p.Glu50, which lies near RAD51D p.Lys297 in the solved structure and could form a complementary charged interaction. Similarly, the RAD51C interfaces were generally missense tolerant, with RAD51D p.Leu164 and p.Ile167 as exceptions, where intolerant variants likely disrupt local helical structure through major physicochemical changes (**Fig. 4e, Extended Data Fig. 8a**). While the Olaparib resistance assay^31^ identified additional sensitive positions such as RAD51D p.Phe176, most intolerant missense variants at this interface require multi-nucleotide substitutions which occur exceptionally rarely in humans, suggesting disruption of RAD51D’s interaction with RAD51C may not be a mechanism contributing to cancer susceptibility.

Lastly, XRCC2 p.Arg114 was tolerant only to charge-preserving substitutions. Since it does not appear to interact with DNA or another BCDX2 or X3CDX2 complex member^23–26^, we hypothesized p.Arg114 may play a critical role for local structural stability. XRCC2 p.Glu75 may form an internal salt bridge with p.Arg114, but tolerance to charge-disrupting variants such as p.Glu75Ala suggests this interaction is unlikely critical for local stability (**Fig. 4d**). However, p.Arg114’s backbone amide forms H-bonds with p.Val78 and p.Cys111, while its sidechain forms an H-bond with the backbone of p.Ile66 (**Fig. 4d**). A similar hydrogen bonding network with p.Val78, p.Cys111, and p.Arg114 was present in the X3CDX2 complex^25,26^ (**Extended Data Fig. 9a**). Consistent with these findings, p.Ile66, p.Val78, and p.Cys111 also did not tolerate substitutions with major physicochemical changes, underscoring the importance of this H-bonding network for XRCC2 stability.

### LOF fitness scores are concordant with pathogenic clinical variant classifications and are enriched in ovarian cancer cases

To understand if SGE-generated fitness scores were concordant with clinical data, we compared *RAD51D* and *XRCC2* functional classifications to ClinVar annotations (**Fig. 5a,b**) (ClinVar release January 2025). Overall, fitness scores discriminated pathogenic/likely pathogenic (PLP) from benign/likely benign (BLB) variants with high accuracy for both genes (AUC=0.994 for *RAD51D* and 1.000 for *XRCC2*; **Fig. 5c,d**). Functional classifications and ClinVar annotations were perfectly concordant for *XRCC2*: all 6 ClinVar PLP variants were LoF, and all 163 BLB variants were functionally normal, resulting in 100% sensitivity and 100% specificity (**Fig. 5b,d**).

**Figure 5.**
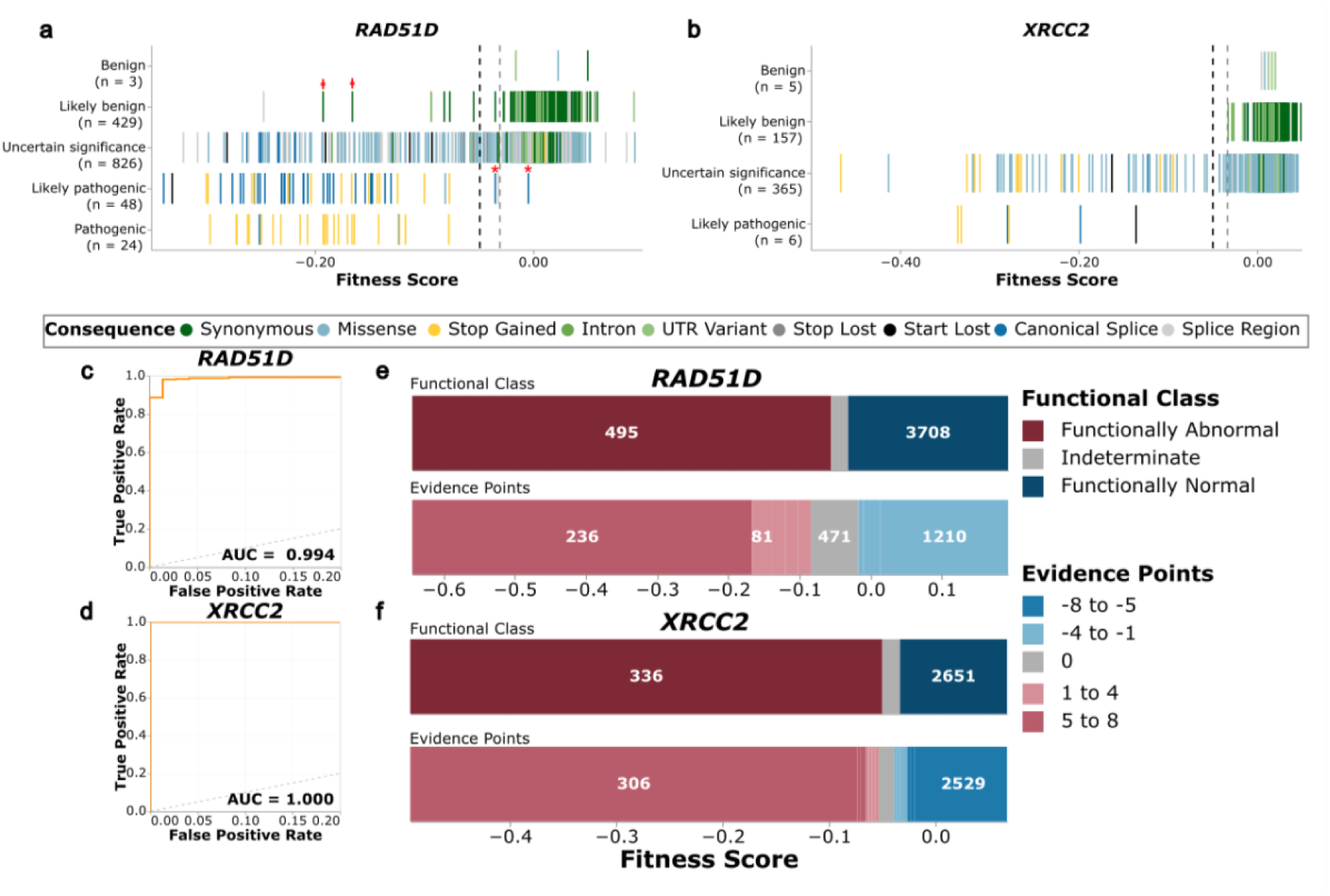
Concordance of SGE functional classifications and ClinVar annotations enables calibration to strong clinical evidence for variant classification. (a) *RAD51D* SNVs (n=1,330) by ClinVar classification (Y-axis) versus fitness score (X-axis). Dashed vertical black and gray lines denote fitness score thresholds for LoF and functionally normal respectively. The molecular consequences of variants are colored as indicated. PLP variants in exon 10 are denoted with red asterisks. BLB synonymous variants that were LoF in SGE with low RNA scores are denoted with red daggers. (b) Analogous to panel (a) for *XRCC2* (n=533). (c) Receiver operator characteristic curve (ROC) for the ability of *RAD51D* fitness scores to discriminate between PLP and BLB variants from ClinVar. (d) Analogous to panel (c) for *XRCC2*. (e) Number of *RAD51D* SNVs (n=4,340) in each functional class (top). Functional categories are colored as indicated. Number of SNVs by ACMG/AMPevidence points received (color) after ExCALIBR^68^ calibration (bottom). (f) Analogous to panel (e) for *XRCC2* (n=3,023).

For *RAD51D*, among 76 ClinVar PLP variants, 72 (94.7%) were LoF by SGE, while 4 (5.3%) were discordant and scored as functionally normal or indeterminate (**Fig. 5a**). Three of the four discordant variants, c.904-2A>C, c.904-2A>T and p.Trp314Ter, occurred in exon 10, which we and others^31^ have demonstrated to be dispensable for RAD51D function (**Fig. 1h**, **Fig. 2a**), suggesting these splice site and stop-gained variants are likely not LoF and merit re-evaluation. The two known likely pathogenic missense variants, p.Met1Lys in exon 1^59^ and p.Ser207Leu in exon 7^64^, were correctly identified as LoF.

Among 433 ClinVar BLB variants in *RAD51D*, 425 (98.2%) had normal fitness by SGE, while 1 (0.2%) were indeterminate and 7 (1.6%) were LoF. All 11 BLB missense variants were correctly classified as functionally normal by SGE. Of the 7 ClinVar BLB variants that were LoF by SGE, five were synonymous changes in exon 3 or exon 4 with low RNA scores (c.225G>A, c.300A>G) or RNA scores close to the low-RNA threshold (c.243C>T, c.321A>G, c.333C>T) (**Fig. 3c, left**); three of these are also predicted to have altered splicing (c.300A>G, c.321A>G, c.333C>T). These data show these BLB synonymous variants can be LoF due to splicing defects and should be reevaluated. The two other BLB variants with LoF fitness scores were intronic (c.263+18G>A) or splice region (c.264-7C>G) variants both in intron 3, thus neither have corroborating RNA scores. Neither are predicted to disrupt splicing, but further exploration of these variants is warranted.

### Functional evidence generated by SGE has high clinical utility

To translate fitness scores into evidence compatible with the ACMG/AMP variant classification guidelines^65–67^, we applied ExCALIBR^68^, which calibrates fitness scores against ClinVar control variants while also taking advantage of gnomAD and synonymous variants to assign evidence points. The distribution of calibrated points was strikingly bimodal, with variants clustering at strongly negative or strongly positive values and relatively few near zero, reflecting the high discriminatory power of the assay (**Fig. 5e,f; Extended Data Fig. 10a,b**). For *RAD51D*, 516 of 4,340 variants were LoF, with 413 (80%) receiving evidence of pathogenicity after calibration, while 470 (10.8%) received 0 points and thus remained uninformative for variant classification (**Fig. 5e**). For *XRCC2*, 336 of 3,023 variants were LoF, with 312 (92.9%) receiving pathogenic evidence after calibration and a similar bimodal distribution of evidence points (**Fig. 5f**). Taken together, calibrated SGE evidence points can be integrated with other lines of variant evidence within the ACMG/AMP framework, providing actionable evidence for the large majority of VUS in both genes.

When calibrated functional evidence is combined with evidence from computational variant effect predictions^69^, 725 *RAD51D* VUS (87%) would receive enough evidence to be reclassified with 634 receiving enough evidence for BLB classification and 91 for PLP. For *XRCC2*, 363 (98%) VUS received enough evidence for reclassification, 312 as BLB and 51 as PLP^39^. Thus, the functional evidence generated by SGE has high clinical utility for classifying variants of *RAD51D* and *XRCC2*.

## Discussion

In this study, we performed saturation genome editing of *RAD51D* and *XRCC2* to generate fitness scores for 5,412 and 3,743 variants, respectively, spanning all coding exons and proximal intronic regions of these two HR genes. By integrating cellular fitness measurements with targeted RNA sequencing, we generated comprehensive functional maps that not only distinguished LoF from functionally normal variants but also revealed molecular mechanisms underlying loss of function. Fitness scores were highly reproducible, concordant with orthogonal HR and protein interaction assays, and recapitulated known structural features of both proteins, including the Walker A and B motifs and other key ATP-coordinating residues. Together, these data provide a near-complete catalog of variant effects for two HR genes implicated in cancer susceptibility and offer mechanistic insight into how individual variants disrupt function at the RNA-and protein-level.

A central finding of this work is the exceptional sensitivity of *RAD51D* to splice-altering variation. Approximately 24% of LoF missense variants in *RAD51D* acted through RNA-mediated rather than protein-mediated mechanisms, a proportion roughly two-fold higher than observed for *BRCA1* and six-fold higher than for *BARD1*, *VHL* or *XRCC2*. This sensitivity appears to be a gene-specific property of *RAD51D*, and we propose that it reflects the transcript architecture of the gene: in HAP1 cells and across human tissues, the full-length *RAD51D* transcript constitutes only <30% of total *RAD51D* expression, with the remainder consisting of non-functional isoforms lacking exon 3 or exons 3–5. Because the full-length transcript represents a minority of total expression, any splice-disrupting variant that further reduces its abundance could be sufficient to cause loss of function. Consistent with this model, exons 3 and 4 harbored the highest concentration of RNA-mediated LoF variants, and RNA scores in these exons explained the majority of the variance in fitness scores for these exons.

Notably, fewer than half of these RNA-mediated LoF variants were predicted by SpliceAI, indicating that current splice prediction tools fail to capture a substantial fraction of potentially clinically relevant splice-disrupting variation. These findings highlight both the value of integrating RNA-level measurements into SGE experimental design and the importance of considering transcript architecture when interpreting variant effects. Direct validation of this transcript architecture model in human carriers, through population-based cohort studies and *in vivo* splicing data, will be important to fully establish its clinical relevance. Specifically, the association of exon 3 and 4 splice-disrupting variants with cancer risk in large population-based cohorts, and comparison of SGE-derived RNA scores with RNA-binding protein interaction data and *in vivo* splicing outcomes, would strengthen the mechanistic and clinical interpretation of this transcript architecture model. These analyses were beyond the scope of the current study but represent a priority for future work.

The functional classes derived from SGE demonstrated high accuracy for distinguishing pathogenic from benign variants. Fitness scores discriminated ClinVar PLP from BLB variants with AUCs of 0.994 for *RAD51D* and 1.000 for *XRCC2*, with perfect concordance for *XRCC2* (100% sensitivity and specificity) and 94.7% sensitivity and 98.2% specificity for *RAD51D*. Importantly, the few discordant variants illuminated rather than undermined the assay’s reliability: three of the four discordant pathogenic variants in *RAD51D* resided in exon 10, which our SGE deletion map and orthogonal data^31^ demonstrate to be dispensable for protein function, suggesting these variants warrant re-evaluation. Similarly, several discordant benign synonymous variants had low RNA scores suggesting potential splice disruption, these too should be re-evaluated. When calibrated against ClinVar controls using the ExCALIBR method, our SGE data provide strong evidence for variant classification. Integration with computational predictors would allow reclassification of 87% of *RAD51D* and 98% of *XRCC2* VUS, providing actionable evidence for the large majority of these uncertain variants.

This work substantially expands the catalog of functional measurements for variants of *RAD51D* and *XRCC2* and demonstrates that comprehensive variant maps can distinguish RNA-level from protein-level mechanisms of loss of function. More broadly, our findings suggest that gene-specific features such as alternative isoform usage and regional NMD sensitivity may shape the landscape of variant effects, and that experimental approaches integrating fitness and RNA-level readouts will be important for variant interpretation across the clinical genome.

## Methods

### RAD51D and XRCC2 essentiality screen in HAP1 cells

To confirm RAD51D and XRCC2 essentiality in HAP1 cells we used a pooled lentiviral CRISPR screen targeting all 10 canonical exons of *RAD51D* (RefSeq:NM_002878.4) and a known alternate exon (RefSeq:NM_001142571.2; chr17:35,116,859-35,117,037) expressed at low frequency in human cells, and all 3 exons of *XRCC2* (RefSeq:NM_005431.2).

For comprehensive exon coverage, all possible gRNAs targeting *RAD51D* and *XRCC2* exons were generated by submitting exonic target sequences and 20-bp flanking intronic sequence to FlashFry^70^ (FlashFry), and filtered for Cutting Frequency Determination (CFD) specificity score of 0.40 or lower^71^. Guide oligos were synthesized by Twist Bioscience with flanking sequences (5’-AGGCACTTGCTCGTACGACGCGTCTCACACCG-3’ and 5’-TTTCGAGACGATGTGGGCCCGGCACCTTAA-3’) for PCR amplification and Golden-Gate cloning into pre-digested LentiGuide Puro-P2A-EGFP (Addgene), gel-purified by excision from a 1% agarose gel. Lentivirus was produced by transfecting eight 10-cm plates of HEK293T cells with the guide library plasmid using Turbofectin 8.0 according to the manufacturer’s protocol (Origene), and viral particles were concentrated using PEG-iT (System Biosciences). To determine an appropriate infectious dose, HAP1-Δ*LIG4*-Cas9-Blast cells were seeded in 12-well plates and transduced in triplicate with 2, 5, 10, or 50 µL of concentrated lentivirus, with transduction efficiency assessed by flow cytometry to determine the volume yielding a multiplicity of infection (MOI) of 0.1. Library representation was confirmed by sequencing on an Illumina NextSeq 2000 platform, with demultiplexing performed using Illumina BCL-Convert software. Twenty-base guide sequences were extracted, and unique guide counts were scored as a proportion of total reads.

For the screen, HAP1-Δ*LIG4*-Cas9 cells were seeded into eight 14-cm plates at 10 million cells per plate and infected the following day at MOI 0.1. Puromycin selection (3 µg/mL) began 30 hours post-infection and was maintained at each passage for the duration of the experiment. Cells were passaged and harvested at days 5, 9, 13, 17, and 21 post-infection. Genomic DNA from days 5, 13, and 17 was extracted using the Qiagen DNeasy Blood & Tissue Kit according to the manufacturer’s protocol. For harvests beyond day 5, which exceeded 20 million cells per replicate, multiple columns were used per biological replicate and eluates were homogenized. Each replicate and timepoint was amplified across 12 KAPA HiFi 2X Mastermix PCR reactions, each containing 500 ng gDNA template, using primers 5’-AGGCACTTGCTCGTACGACG-3’ and 5’-TTAAGGTGCCGGGCCCACAT-3’. Reactions from each replicate and timepoint were pooled prior to AMPure XP bead cleanup and indexing with Illumina adapters and sample-specific indexes. Sequencing and scoring were performed as described above, and the log2 fold change (log2FC) of guide scores relative to the day 5 library were plotted for all three timepoints (**Extended Data Fig. 1a,d**). Guides producing significant depletion were selected for use in full SGE experiments.

### sgRNA design and cloning for saturating genome editing

For SGE experiments, sgRNAs were designed using the CRISPR design function in Benchling, with off-targetCFD scores and on-target scores calculated for NGG PAM sites with 20-nt guides against GRCh38 (hg38), using the scoring algorithm of Doench et al^71^. Guides were selected based on the following criteria: minimum predicted off-target activity, maximum predicted on-target efficiency, amenability to introduction of a synonymous change at the protospacer or PAM site to prevent Cas9 recutting following HDR, and strong depletion scores in the pooled CRISPR screen. For PAM site modifications, C>G or G>C transversions were preferred. When no guide had a PAM site permitting a synonymous change, two synonymous changes were instead introduced within the protospacer sequence. All sgRNAs were synthesized by Integrated DNA Technologies (IDT) and cloned as previously described (protocols.io: dx.doi.org/10.17504/protocols.io.3byl46ddjgo5/v1). Cloned constructs were confirmed by Sanger sequencing, and sequence-verified plasmids were propagated overnight in 150 mL LB broth and purified using the ZymoPURE II Maxiprep kit with EndoZero spin columns (Zymo Research).

### HDR library design and cloning

The SGE target sequences (∼120 bp) were retrieved from the hg38 genome assembly based on MANE select transcripts (RefSeq:NM_002878.4 for *RAD51D* and NM_005431.2 for *XRCC2*), and the required synonymous substitutions for the selected sgRNAs were introduced into the reference sequences. Where possible, a second synonymous substitution was introduced at an alternative CRISPR cut site; if not feasible, it was placed on the opposite side of the target or within a flanking intronic region.

Pooled oligonucleotides were array-synthesized by Twist Bioscience. Target-specific oligonucleotides were amplified from the pool using target-specific primers (**Supplementary File 3**) and purified using Sera-Mag Select beads (Cytiva). All subsequent steps, including homology arm cloning, HDR library assembly, transformation, and purification, were performed as previously described^37^.

### Cell culture, transfection, and time point sampling

HAP1-Δ*LIG4*-Cas9 cells (Horizon Discovery) were cultured in Iscove’s Modified Dulbecco’s Medium (IMDM, Gibco) supplemented with 10% fetal bovine serum (MilliporeSigma) and routinely sorted for the 1N haploid population prior to use. All SGE experiments were performed as previously described^37^, with the following exceptions. For *RAD51D*, 10 transfections were performed per SGE region (1 negative control and 9 replicates); for *XRCC2*, 7 transfections were performed per region (1 negative control and 6 replicates). Sequencing timepoints also differed by gene: for *RAD51D*, samples from days 5, 13, and 17 were used for sequencing, whereas for *XRCC2*, samples from days 5 and 13 were used.

### gDNA and RNA extraction and sequencing preparation

gDNA extraction, quantification, and sequencing library preparation were performed as previously described^37^, with the exception that for *RAD51D*, gDNA was additionally extracted from day 17 cell pellets using a DNeasy kit (Qiagen) and quantified using the Qubit 1x Broad Range Kit (Invitrogen), and day 17 samples were included in sequencing library preparation following the same protocol. Gene-specific primers used for gDNA library preparation for both *RAD51D* and *XRCC2* are provided in **Supplementary File 3**.

RNA sequencing libraries were prepared from day 5 samples for both *RAD51D* and *XRCC2*. Reverse transcription was primed using gene-specific 3’ UTR primers, and all subsequent PCR steps used gene-specific primers targeting flanking exons of each region of interest (**Supplementary File 3**). All remaining library preparation steps were performed as previously described^37^.

### Processing of sequencing data

Paired-end reads were adapter-trimmed and merged with SeqPrep (version 1.3.2) with the following options:-A GGTTTGGAGCGAGATTGATAAAGT-B CTGAGCTCTCTCACAGCCATTTAG-M 0.1-m 0.001-q 20-o 20. Reads containing N bases were discarded. Merged reads were mapped to target-specific, GRCh38 reference sequences using the mem algorithm in bwa (version 0.7.17-r1188) to produce target-specific BAM files.

Counts of each programmed SNV within each replicate and at each timepoint (henceforth, “dataset”), and within the repair template library, were extracted from each BAM file by a custom Python script. All mismatches between the merged read sequence and the reference sequence were identified. Reads were considered valid if they had the expected length, contained all fixed edits, and had a single additional mismatch within the edited region of the target, which was taken as the programmed edit. Additional mismatches within the merged read but falling outside of the edited region were tolerated. Filtering was relaxed for SGE targets containing homopolymer runs with length ≥4. In such cases, a change in sequenced homopolymer length of up to two was allowed.

Programmed variants with fewer than ten observations in the repair template library or any dataset were excluded from analysis. Counts of each programmed 3-bp deletion within each dataset, and within the repair template library, were similarly extracted from BAM files by a custom Python script. CIGAR strings were parsed to identify alignments containing a single 3-bp deletion in the merged read sequence. Reads containing such deletions were further examined to ensure that all fixed edits were present, except for those that overlapped with the identified, programmed deletion. Reads containing sequence-level mismatches within the edited region as well as a single programmed deletion were discarded; sequence mismatches outside of the edited region were tolerated. Each variant’s count within each dataset was converted to a variant frequency by dividing the integer count by the number of valid reads within the dataset. The editing rate per dataset was calculated by dividing the sum of valid SNV-containing reads and valid deletion-containing reads by the total number of reads.

### Smoothing

A frequency ratio was computed for each variant in each day 5 replicate, comprising the log_2_ of its frequency at day 5 over its frequency in the repair template library. A one-dimensional LOESS smoother, implemented in Python with the loess package (version 2.1.2) with the option span=0.20, was used to attenuate the presumed position effect on the observed variant counts owing to the location of the cut-site relative to the programmed edit. The smoother was fit separately within each replicate to the log2 ratios at day 5 to yield fitted log_2_ ratios. These positional fits were then subtracted from the day 5 log_2_ frequency ratios, and from each replicate’s cognate day 13 and (for *RAD51D*) day 17 log_2_ frequency ratios, to yield adjusted log_2_ ratios. The model was fit only on those variants whose frequencies at day 5 were no less than half of their starting frequencies.

### Scoring

A continuous-time linear regression model, implemented in the linregress module of scipy (version 1.14.0), was used to estimate SNV and deletion scores as a function of the adjusted log2 frequency ratios within each replicate and each timepoint: 0, 5, and 13 days for both genes, plus an additional day 17 timepoint for *RAD51D*. The log_2_ variant frequency at the starting timepoint was set to 0. Estimated log_2_ fold-changes per unit time (i.e., per day) were taken as the fitness score for each variant. For variants covered by two adjacent SGE targets, frequencies from both targets were jointly modeled to produce a single score.

RNA scores were generated by calculating the log_2_ frequency ratio of each variant in the RNA to its respective day 5 DNA frequency in the corresponding replicate. Since the genomic region sequenced in DNA samples is larger than the region sequenced in RNA samples, prior to RNA scoring, day 5 DNA frequencies were adjusted to include only positions sequenced in RNA samples. RNA scores were independently calculated for each experimental replicate and collapsed into a final score by taking the median of all replicate scores. For variants covered by two adjacent SGE targets, the final reported score is the mean of both individual scores.

### Variant annotation

The expected molecular consequence(s), HGVS identifiers, and amino acid change of each programmed variant were annotated by the Ensembl Variant Effect Predictor (vep), version 115, using the MANE Select transcripts for *RAD51D* and *XRCC2*. For variants with multiple predicted molecular consequences, only the consequence with the highest predicted impact was retained.

### Functional class and RNA expression class assignment

A two-component Gaussian mixture model, implemented in scikit-learn (version 1.7.1), was fit to the SNV scores. A separate model was fit for each of the two genes. The model’s initial component means were set to the mean of the middle 95% of the score distribution of expected-neutral SNVs—specifically, synonymous and intronic SNVs—and the mean score of all stop-gained SNVs outside of exons 1 and 10 (for *RAD51D)* or outside of exon 1 (for *XRCC2)*. The probability of belonging to each component was estimated for each SNV, and those whose probability of assignment to a component exceeded 0.95 were assigned that component’s label: “functionally abnormal” and “functionally normal” for the components with the lesser and greater means, respectively. Remaining SNVs were labeled “indeterminate.” The approximate score threshold values for functional class assignment were identified by taking the scores of the variants that minimized the distance to one of the 0.95 probability cutoffs. These two score thresholds were then applied to the 3-bp deletion scores, without refitting, to yield the same three functional classes. The final fitness score thresholds for the functionally abnormal and functionally normal classes were-0.0487 and-0.0304 (for *RAD51D*), and-0.0498 and-0.0335 (for *XRCC2*).

RNA scores were discretized into “low” and “normal” RNA expression classes using the mean and standard deviation of the synonymous variant RNA score distribution. The RNA threshold distinguishing low from normal RNA was set at the mean minus 3 standard deviations of this distribution. The RNA score thresholds for low and normal RNA were −2.7856 for *RAD51D* and-0.6603 for *XRCC2* (**Supplementary File 1, 2)**.

Stop-gained variants were considered NMD-susceptible when located outside the first exon or more than 100bp from the ATG, outside the last exon and more than 50-55bp upstream the last exon-exon junction, consistent with established NMD rules^72,73^.

### Variant filtering

Programmed SNVs in the same codon as a suspected cell line SNP or a fixed edit required for SGE library design were scored; however, variant annotations based on the reference *RAD51D* or *XRCC2* transcript may assign incorrect molecular consequences to these altered codon contexts. Consequently, these variants were removed from the final data set.

### Long-read RNA-seq

mRNA was isolated using the Qiagen RNeasy Mini Kit (74104) and subjected to polyA-mediated cDNA synthesis as previously described^74^. cDNA was then concatemerized using the Kinnex Full-Length RNA kit (PacBio, 103-072-000). Libraries were sequenced on the Revio platform using the SMRT-Nx Revio Chemistry.

### Calibration and comparison to ClinVar

SGE functional classifications were compared to clinical variant interpretations using the January 2025 ClinVar release. Variants with review status “criteria provided, single submitter” or higher were included and variants with “no assertion criteria provided” were excluded. Variants were grouped as pathogenic/likely pathogenic (PLP) or benign/likely benign (BLB), with conflicting interpretations and variants of uncertain significance excluded from the analysis. SGE assay performance was evaluated using receiver operating characteristic curve (ROC) analysis, and concordance was assessed by calculating sensitivity as the proportion of PLP variants correctly identified as loss-of-function and specificity as the proportion of BLB variants correctly identified as functionally normal.

To translate SGE fitness scores into standardized evidence compatible with ACMG/AMP variant classification guidelines, we applied ExCALIBR^68^. ExCALIBR jointly models score distributions of PLP, BLB, synonymous, and gnomAD v4.1 population variants and estimates gene-specific prior probabilities of pathogenicity to compute variant-specific positive likelihood ratios. The assigned posterior probabilities of pathogenicity are translated into evidence compatible with ACMG/AMP guidelines. For *RAD51D* and *XRCC2* SGE fitness scores, we used 3-component and 2-component ExCALIBR models, respectively, with all other parameters default. To assess VUS reclassification potential, calibrated functional evidence was combined with computational variant effect prediction evidence as described by Tejura et al^39^.

## Resource Availability

### Lead contact

Further information and requests for resources and reagents should be directed to and will be fulfilled by the lead contacts, Silvia Casadei and Lea M. Starita

### Materials availability

SGE assay standards used for fitness score calculation are available through the IGVF Consortium in the IGVF Assay Standards at: https://docs.google.com/document/d/1bVftuEWZAupaBPmSEErPYfw6xv4xdgPn/edit

Please contact the corresponding authors with requests for any further reagents generated by this study.

### Data and code availability

All software and supporting files are available at https://github.com/bbi-lab/sge-pipeline/.

Code and data to regenerate all figures and analyses from supplementary files are available at: https://github.com/bbi-lab/RAD51D_XRCC2_analysis

Raw sequencing reads can be downloaded from IGVF (https://data.igvf.org/): RAD51D: IGVFDS8628CRZA; XRCC2: IGVFDS2228GRCP.

SGE scores have been deposited in MaveDB. RAD51D: urn:mavedb:00001260-a-2; XRCC2: urn:mavedb:00001264-a-2.

## Acknowledgements

We would like to thank current and former members of the Starita lab, Brotman Baty Advanced Technology Lab, IGVF Coding Variant Focus Group and ClinGen / AVE Functional Data Working Group for helpful discussions. The AI Claude (Anthropic) was used to refine text. This work was supported by the following awards: National Institute of Health (NIH) National Human Genome Research Institute (NHGRI) Impact of Genomic Variation on Function (IGVF) consortium center awards: the Center for Actionable Variant Analysis (CAVA to D.M.F. and L.M.S (UM1HG011969) and the Radivojac Center to P.R. (U01HG012022). An NIH NHGRI grant supported R.K.G (1RM1HG010461). Additional financial support was provided by an NHGRI Advancing Genome Medicine Research consortium award (R01HG013025, L.M.S. and D.M.F.), an NHGRI GREGoR consortium award (U01HG011744, support to L.M.S) and support from Google.org to A.B.S. Further funding supported N.J.G. (5T32ES019851-10) and J.O.K. (R35 GM153286). R.K.G acknowledges support from a Washington Research Foundation postdoctoral fellowship. L.M.S. acknowledges support from the Brotman Baty Institute for Precision Medicine.

## Author contributions

S.C., D.M.F., and L.M.S. conceptualized the project. Methodology was developed by S.C., M.W.S., R.K.G., P.R., D.M.F., and L.M.S. Software was developed by M.W.S., R.K.G., and R.S. Validation was performed by S.C., R.S., N.J.G., J.B., D.M.F., and L.M.S. Formal analysis was carried out by S.C., M.W.S., I.W., N.S., R.K.G., J.B., P.G., D.M.F., and L.M.S. Data curation was performed by R.K.G. and M.T. Investigation was conducted by S.C., M.S., I.W., N.S., S.B., R.K.G., L.R.S., C.W., O.S., A.H., A.X., J.R., D.M.F., and L.M.S. Resources were provided by S.C., M.W.S., I.W., N.S., S.B., R.K.G., L.R.S., C.W., O.S., A.H., A.X., J.O.K., D.M.F., and L.M.S. The original draft was written by S.C., I.W., R.K.G., D.M.F., and L.M.S. Review and editing were performed by S.C., I.W., R.K.G., N.J.G., K.A.B., J.O.K., D.M.F., and L.M.S. Visualization was carried out by S.C., M.W.S., I.W., R.K.G., R.S., P.R., D.M.F., and L.M.S. Supervision was provided by S.C., A.B.S., P.R., D.M.F., and L.M.S. Project administration was handled by S.C., S.H., L.M., D.M.F., and L.M.S. Funding acquisition was led by A.B.S., P.R., D.M.F., and L.M.S.

## Conflicts of Interest

Jacob O. Kitzman serves on the scientific advisory board of Myome, Inc. and Douglas M. Fowler is a member of the Alloz Bio scientific advisory board

## Extended Data Figures

**Extended Data Figure 1.**
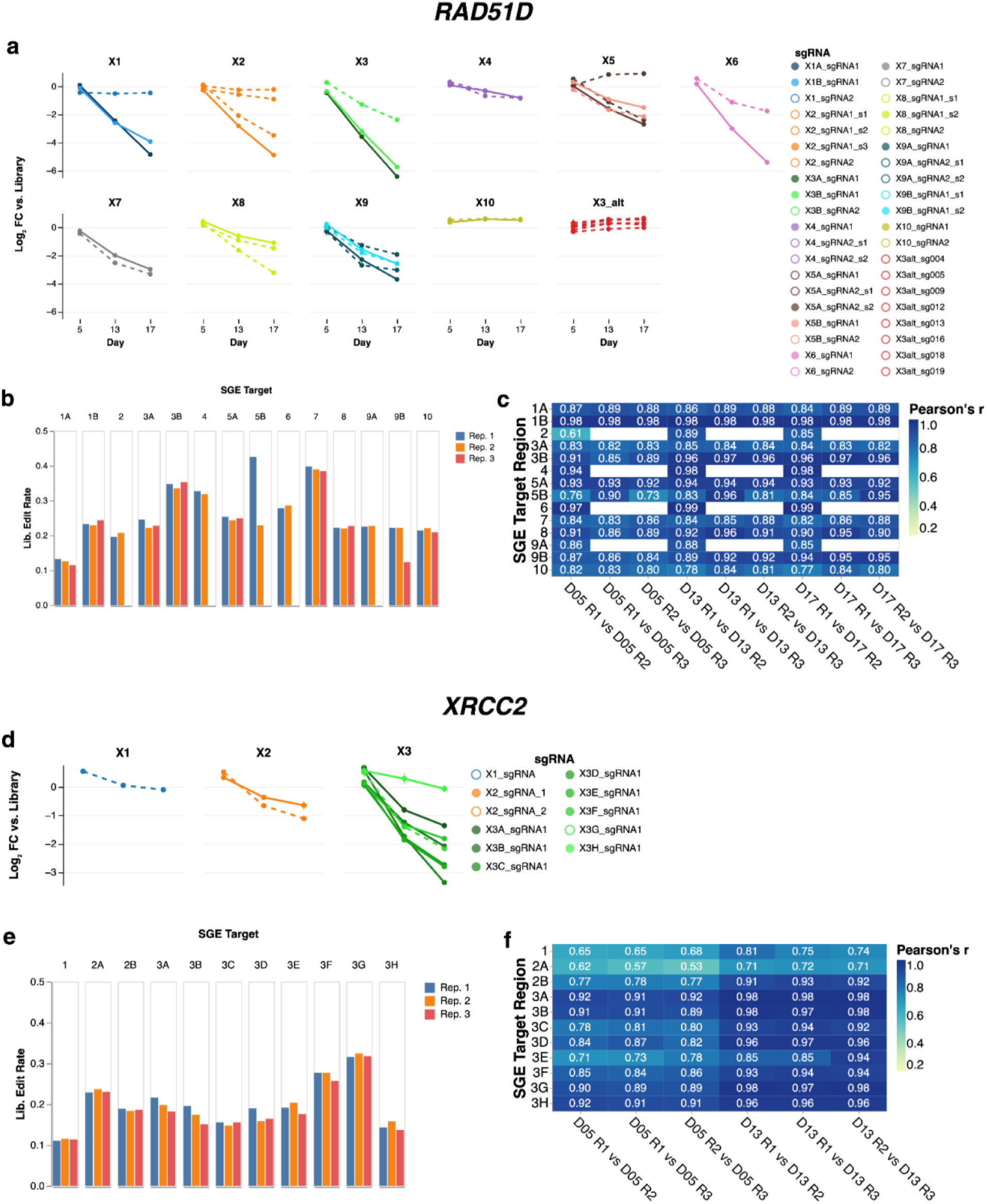
Selection of sgRNAs and quality control of *RAD51D* and *XRCC2* SGE targets. (a) Log2-fold change (Y-axis) of counts of sgRNAs tested in CRISPR-Cas9 screen of *RAD51D* exons over three timepoints (X-axis). sgRNAs are colored as indicated. Solid and dotted lines denote guides used in tissue culture editing experiments or guides used only in the screen respectively. “X3_alt” denotes guides targeting the alternate exon 3 of *RAD51D*. (b) Editing rates (Y-axis) generating usable reads across QC-passing targets in *RAD51D* (X-axis) and replicates (bar color). Usable reads are defined as having all fixed edits and a single programmed 3-bp deletion. (c) Heat map of Pearson’s correlation between replicates (X-axis) for QC-passing targets (Y-axis) in *RAD51D*. Pearson’s correlation is colored for each target and replicate comparison. (d) Analogous to panel (a) for *XRCC2*. (e) Analogous to panel (b) for *XRCC2*. (f) Analogous to panel (c) for *XRCC2*.

**Extended Data Figure 2.**
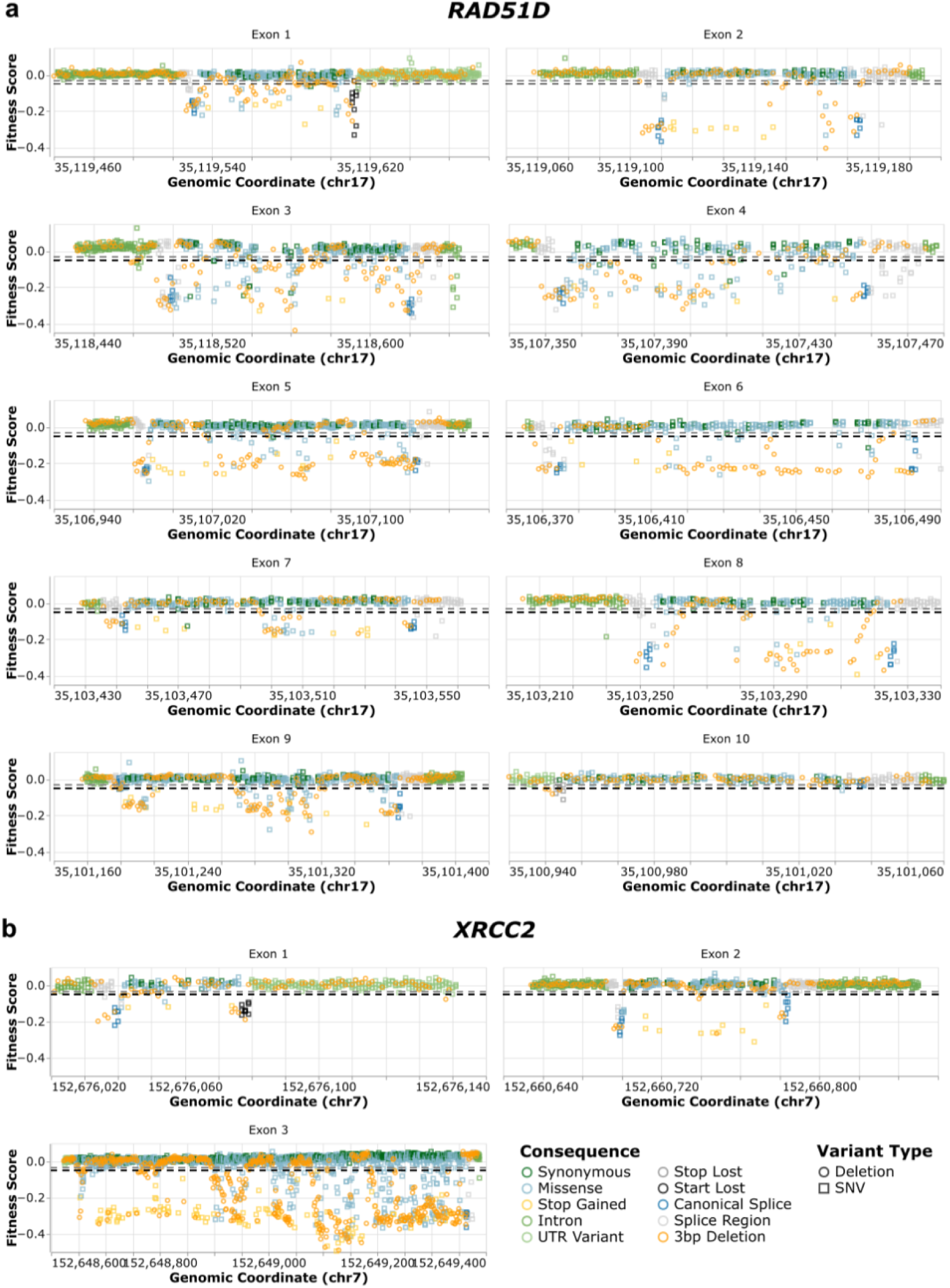
Fitness scores for programmed SNVs and 3-bp deletions by variant class and exon. (a) Fitness scores for all *RAD51D* variants plotted by genomic coordinate (chr17) for each of the 10 coding exons. (b) Fitness scores for all *XRCC2* variants plotted by genomic coordinate (chr7) for exons 1-3. Each point represents a single variant, colored by mutational consequence and shaped by variant type (circle: 3-bp deletion; square: SNV), as indicated. Dashed horizontal gray and black lines indicate the functionally normal and functionally abnormal classification thresholds. Genomic coordinates are based on GRCh38/hg38.

**Extended Data Figure 3.**
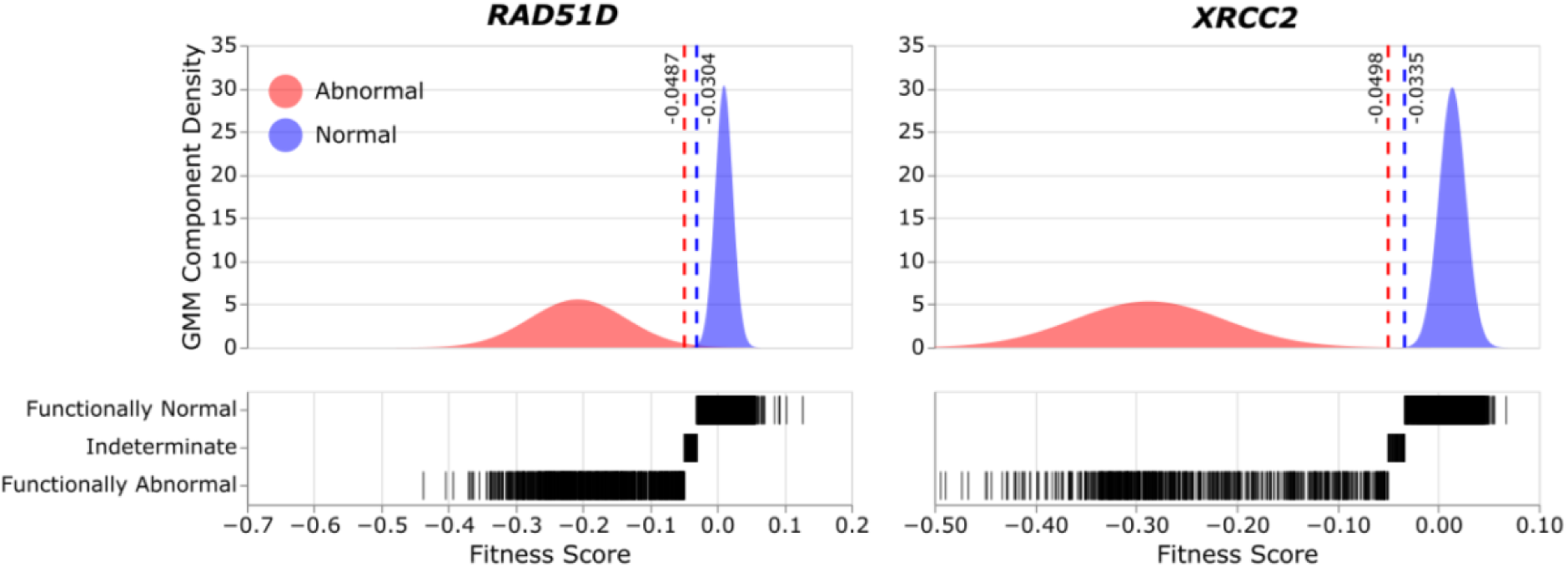
Gaussian mixture model-based functional class assignment for *RAD51D* and *XRCC2*. For each gene, a Gaussian Mixture Model (GMM) was fit to the distribution of SNV fitness scores (**Methods**). Estimated densities of the two components after fitting are shown in red and blue (*top*). Dashed vertical lines indicate the score threshold values separating the three functional classes. Each SNV and 3-bp deletion fitness score was compared to the threshold values to produce functional class assignments (*bottom*).

**Extended Data Figure 4.**
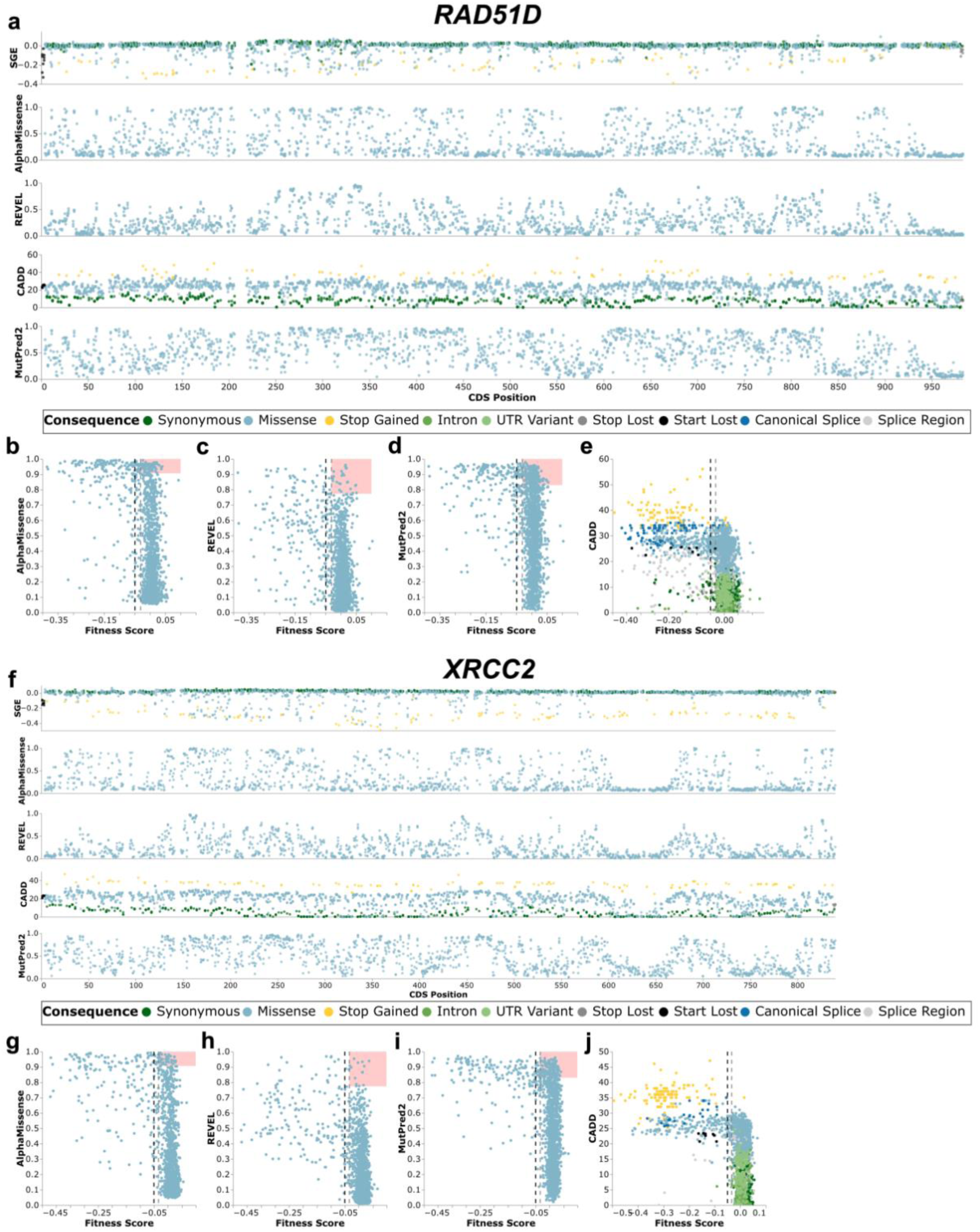
Comparison of SGE fitness scores to computational predictions of variant effects. (a) SGE fitness scores for all *RAD51D* variants plotted by CDS position. Variants are colored by predicted consequence: missense (light blue), synonymous (dark green), stop-gained (yellow), intronic (olive green), UTR (light green), stop-lost (grey), start-lost (black), splice site (dark blue), and splicing variants (light grey). (b) - (e) Comparison of fitness scores and computational variant effect predictors for *RAD51D*: scatterplots of AlphaMissense (b), REVEL scores (c), and MutPred2 (d) scores against fitness scores for missense variants. The vertical dashed line indicates the SGE functional classification threshold; the pink shaded region highlights variants predicted to have at least moderate evidence toward pathogenic classification^43,75^ that are functionally normal by SGE. Among missense variants with normal cellular fitness, MutPred2 showed the highest rate of pathogenic calls (12.1%) followed by AlphaMissense (4.2%) and REVEL (1.6%). (e) CADD scores plotted against fitness scores for all variant consequences. Vertical dashed lines indicate the SGE functional classification threshold. (f) SGE fitness scores for all *XRCC2* variants plotted by CDS position.Color scheme as described in (a). (g) - (j) Comparison of fitness scores and computational variant effect predictors for *XRCC2*: scatterplots of AlphaMissense (g), REVEL (h), and MutPred2 (i) scores against fitness scores for missense variants. The vertical dashed line indicates the SGE functional classification threshold; the pink shaded region highlights variants predicted pathogenic that are functionally normal by SGE. Among missense variants with normal cellular fitness, MutPred2 showed the highest rate of pathogenic calls (8.3%) followed by AlphaMissense (3.0%) and REVEL (1.2%). (j) CADD scores plotted against fitness scores for all variant consequences. The vertical dashed line indicates the SGE functional classification threshold.

**Extended Data Figure 5.**
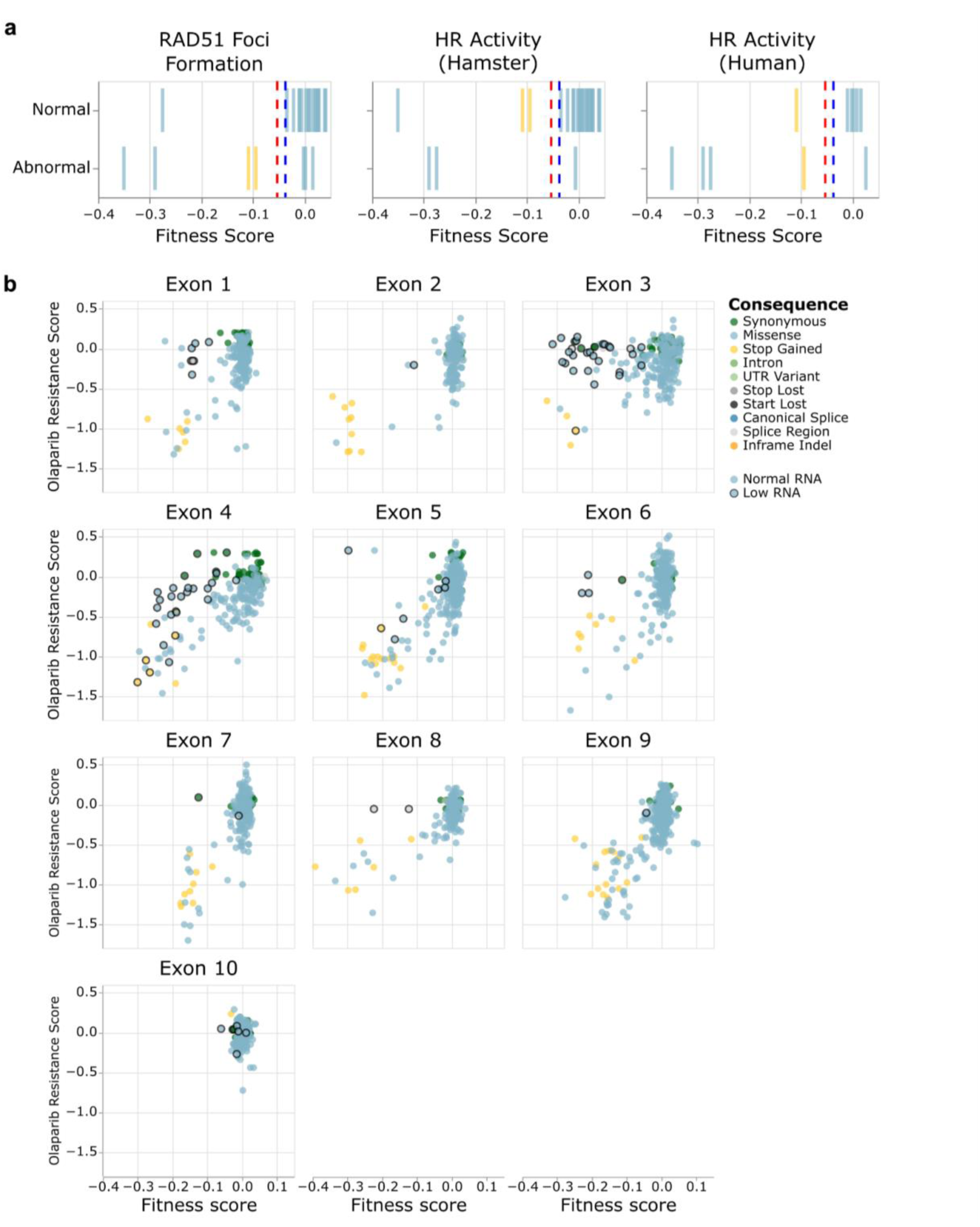
Concordance between *RAD51D* and *XRCC2* fitness scores and orthogonal assays. (a) Comparison of *XRCC2* fitness scores and functional classification from three independent, orthogonal assays^47^: RAD51 foci formation in *irs1* hamster cells (*left)*, homologous recombination (HR) activity in *irs1* hamster cells (*middle*), and HR activity in HEK293 cells (*right*). Vertical dashed lanes indicate the two SGE functional classification thresholds. (b) Comparison of *RAD51D* fitness scores to Olaparib resistance scores^31^.

**Extended Data Figure 6.**
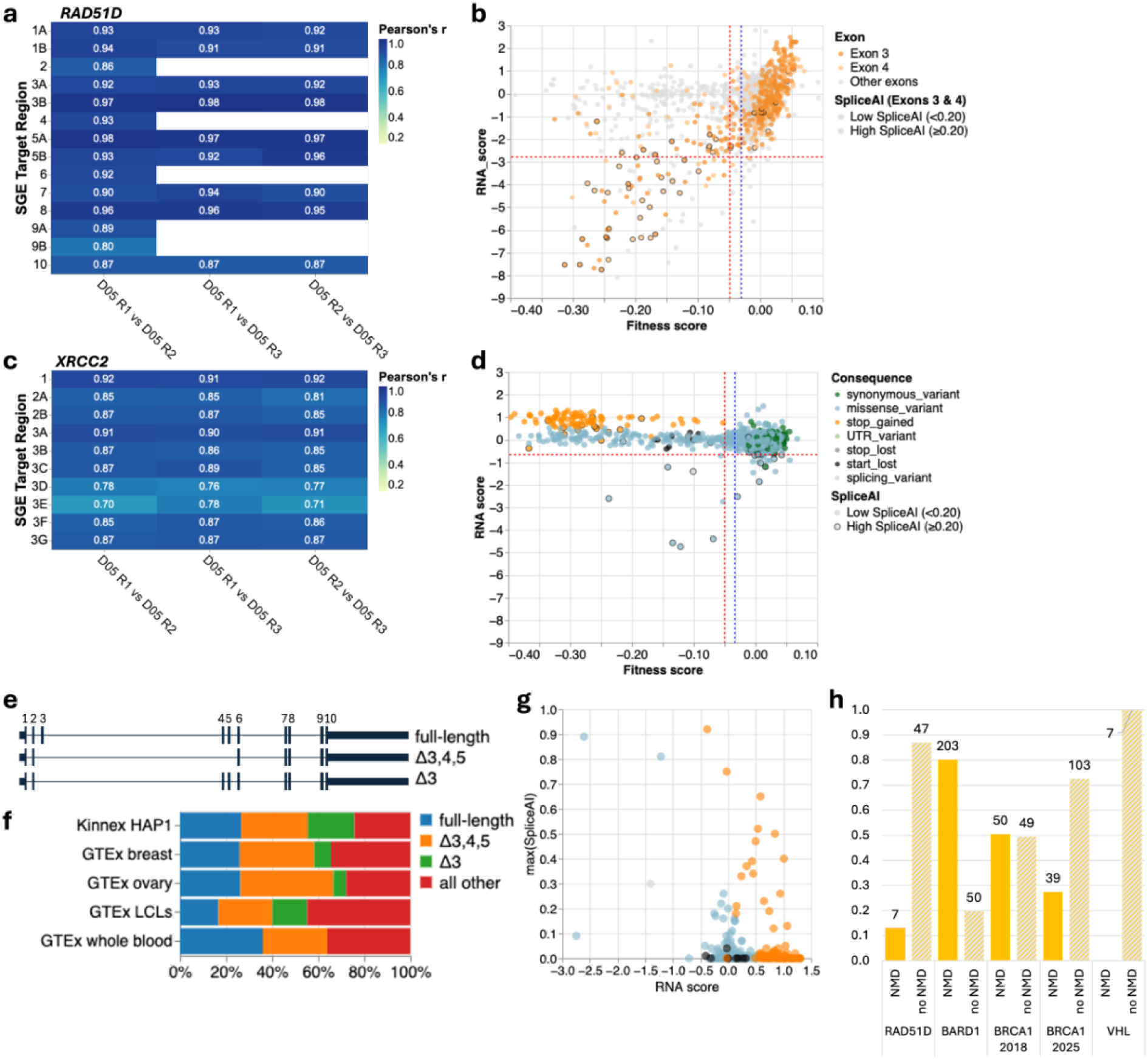
RNA scores, transcript isoforms, and NMD landscape for *RAD51D* and *XRCC2*. (a) Heat map of Pearson’s correlation between RNA replicates (X-axis) collected at day 5 (D05) for each QC-passing target (Y-axis) in *RAD51D*. Pearson’s correlation is colored for each target and replicate comparison. (b) Scatterplot of RNA scores versus SGE fitness scores for *RAD51D* exons 3 and 4 with SpliceAI score overlaid as outline. Variants in all other exons are shown in grey. For exons 3 and 4, outlined circles indicate variants with high SpliceAI max score (≥0.20); open circles indicate variants with low SpliceAI max score (<0.20). (c) Analogous to panel (a) for *XRCC2*. (d) Scatterplot of RNA scores versus SGE fitness scores for *XRCC2* coding variants colored by consequence. The red dashed horizontal line indicates the low RNA threshold (−0.6603) at three standard deviations below the synonymous variant mean. The blue and red dashed vertical lines indicate the SGE thresholds for normal (−0.0335) and LoF (−0.0498) fitness scores, respectively. Outlined circles indicate variants with high SpliceAI max score (≥0.20); open circles indicate variants with low SpliceAI max score (<0.20). (e) Schematic of three predominant *RAD51D* transcript isoforms. Full-length transcript (top), Δ3,4,5 (middle) and Δ3 isoform are shown, with exons represented as filled blocks and introns as connecting lines. (f) Proportion of *RAD51D* transcript isoforms across cell types and tissue. Percentages of full-length, Δ3,4,5 and Δ3 isoforms for HAP1 obtained by long-read RNA sequencing (Kinnex, PacBio), and for breast, ovary, lymphoblastoid cells (LCLs) and whole blood, with the latter four derived from GTEx expression data^49,50^. (g) Scatterplot of XRCC2 RNA scores (X-axis) and SpliceAI maximum scores (Y-axis) for functionally abnormal *XRCC2* variants only. Colors indicate variant consequences as described in panel b. (h) Among stop-gained variants predicted to be susceptible to canonical NMD, the proportion that show low RNA consistent with NMD (NMD, yellow solid bars) versus those that escape (no NMD, hatched bars) shown for *RAD51D* and published SGE datasets from *BARD1*^37^, *BRCA1*^33,34^, and *VHL*^41^. *XRCC2* is not shown as none of its 144 stop-gained variants are predicted to be NMD-susceptible, consistent with its three exon structure. Numbers above bars indicate variant counts.

**Extended Data Figure 7.**
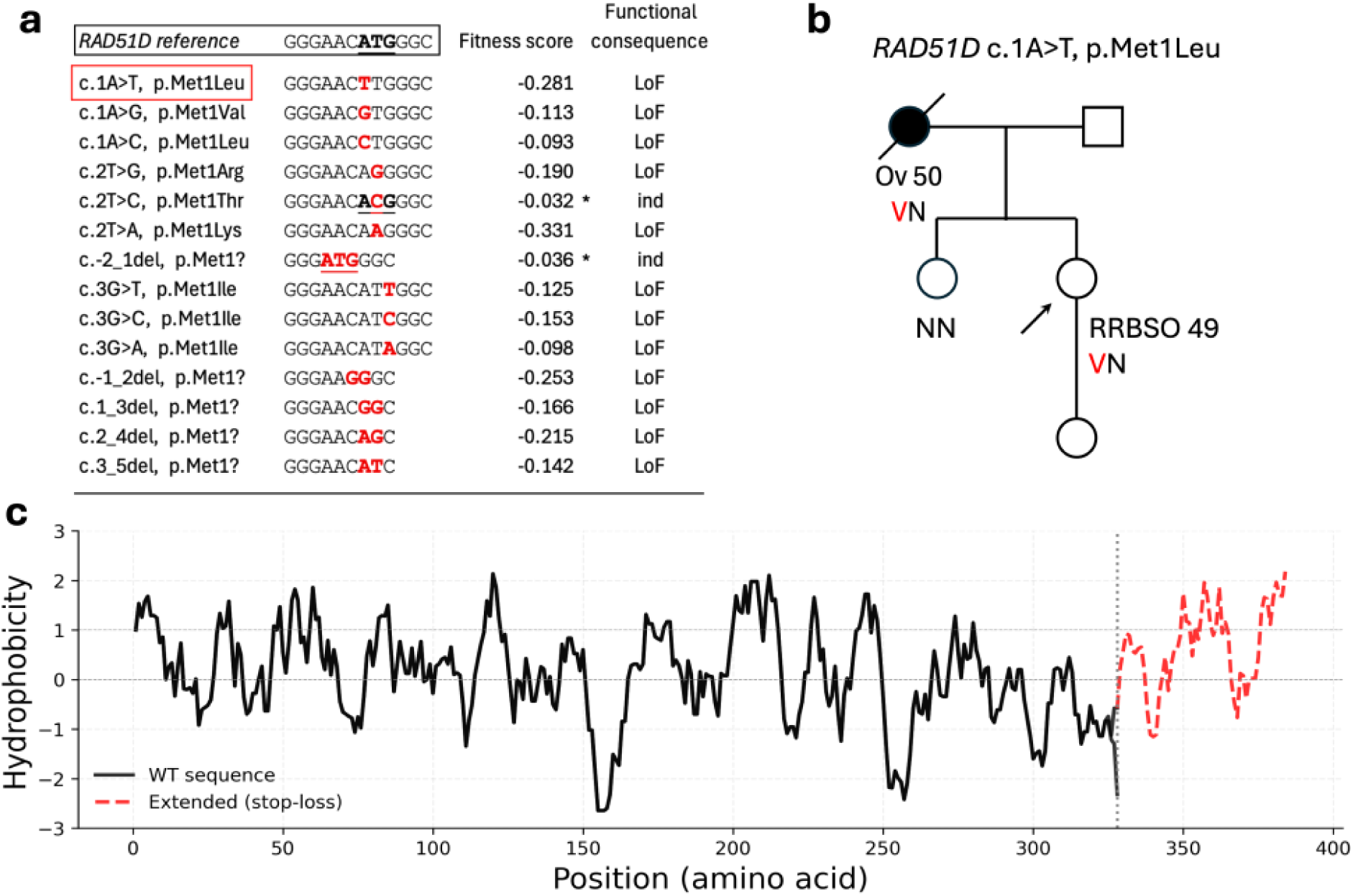
start-and stop-lost variants reveal distinct mechanisms by which translation can be preserved or ablated. (a) Functional impact of *RAD51D* start-loss variants. The *RAD51D* reference sequence with strong Kozak consensus (positions-3,-1, +4: A, C, G) is shown at top. Two functionally indeterminate variants (marked with asterisks) demonstrate compensatory mechanisms: c.-2_1delACA maintains methionine while disrupting the Kozak sequence, and c.2T>C (p.Met1Thr) creates a non-canonical ACG start codon (ATG > ACG) that support translation via Met-tRNAi, incorporating methionine rather than threonine^56^. The variant c.1A>T (p.Met1Leu) shown in the pedigree in **panel b** is indicated with a red frame. Variant nucleotides are highlighted in red. (b) Family pedigree for RAD51D c.1A>T (p.Met1Leu). The proband’s mother, who carries the variant, had endometrioid ovarian cancer (serous) at age 50 and died at age 54. The proband (arrow) carries the variant and underwent risk-reducing bilateral salpingo-oophorectomy at age 49 (RRBSO 49). Her sister does not carry the variant and has not had RRBSO. (c) Hydrophobicity profile of *RAD51D* wild-type and stop-loss extension. Stop-loss variants introduce a 168 bp 3’ UTR sequence predicting a 56-amino-acid extension (positions 329-384, red dashed line) that avoids non-stop decay. The extension is predominantly hydrophobic (56% overall, 73% in the final 15 residues) with hydrophobicity increasing towards the C-terminus. The vertical dotted line indicates the normal stop codon at position 329. The horizontal dashed line indicates the moderately hydrophobic threshold (>1.0, Kyte-Doolittle scale^76^.

**Extended Data Figure 8.**
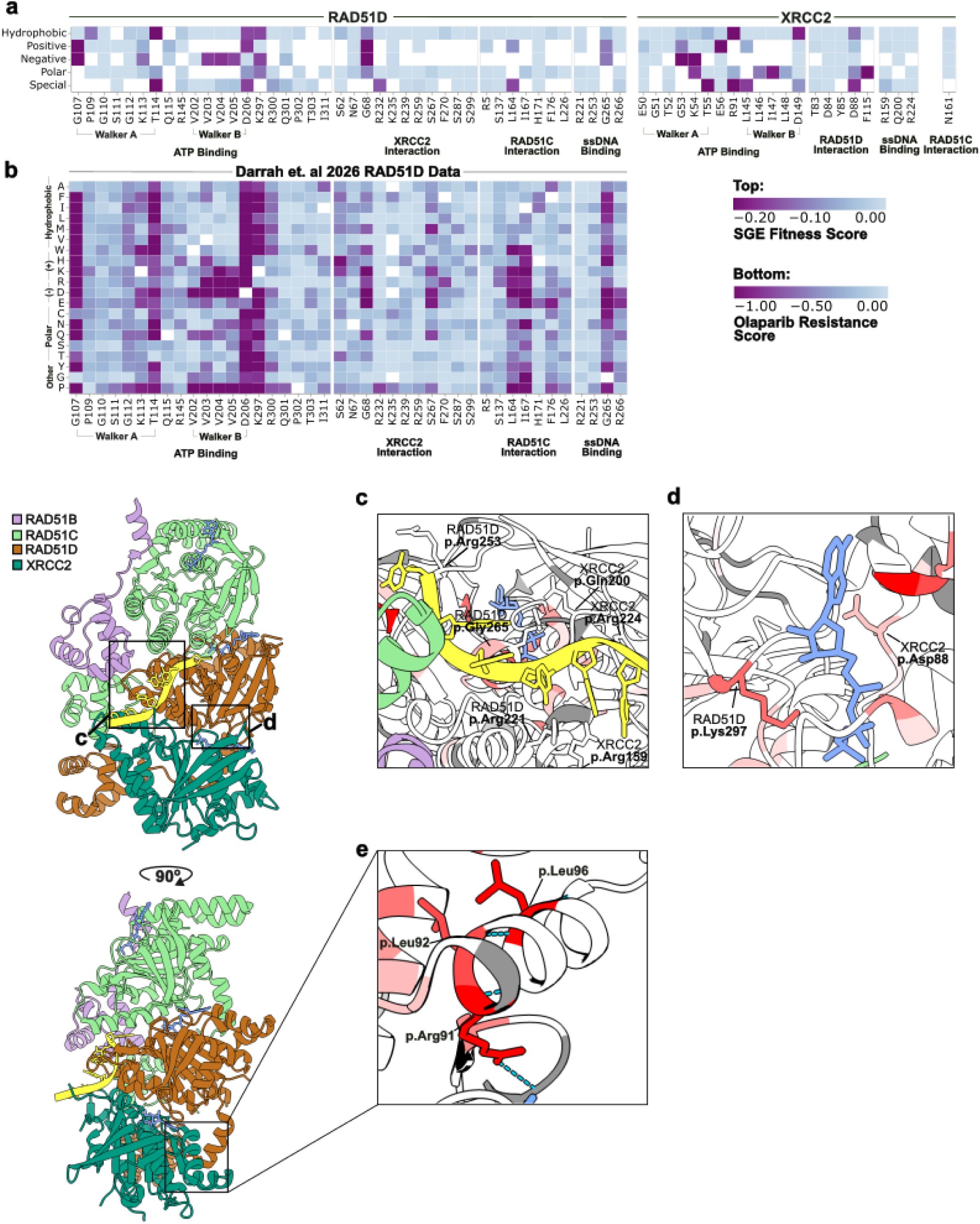
Structural and functional consequences of missense variants on RAD51D and XRCC2. (a) Heatmap of residues previously reported^23,24^ in RAD51D (left) and XRCC2 (right) biochemical interactions. Y-axis denotes properties of substituted amino acids. X-axis denotes original amino acid and residue position. Fitness scores are colored as indicated. For residues where multiple substitutions introduced the same biochemical property, the median fitness score was calculated. (b) Heatmap of residues previously reported^23,24^ in RAD51D biochemical interactions. Y-axis denotes substituted amino acids. X-axis denotes original amino acid and residue position. Residues are grouped by their biochemical function. Functional scores measured by Darrah et al. 2026^31^ are colored as indicated. (c) ssDNA interaction interface of RAD51D and XRCC2. Residues reported to interact(PDB: 8GBJ^23^) are highlighted. Residues are colored by median fitness score. Residues not scored are colored in gray. (d) RAD51D p.Lys297 and XRCC2 p.Asp88 at the RAD51D-XRCC2 interface (PDB: 8GBJ^23^). Residues are colored by median fitness score. (e) XRCC2 p.Arg91 and hydrogen bonding interactions likely required for the stability of XRCC2(PDB: 8GBJ^23^). Residues are colored by median fitness score.

**Extended Data Figure 9.**
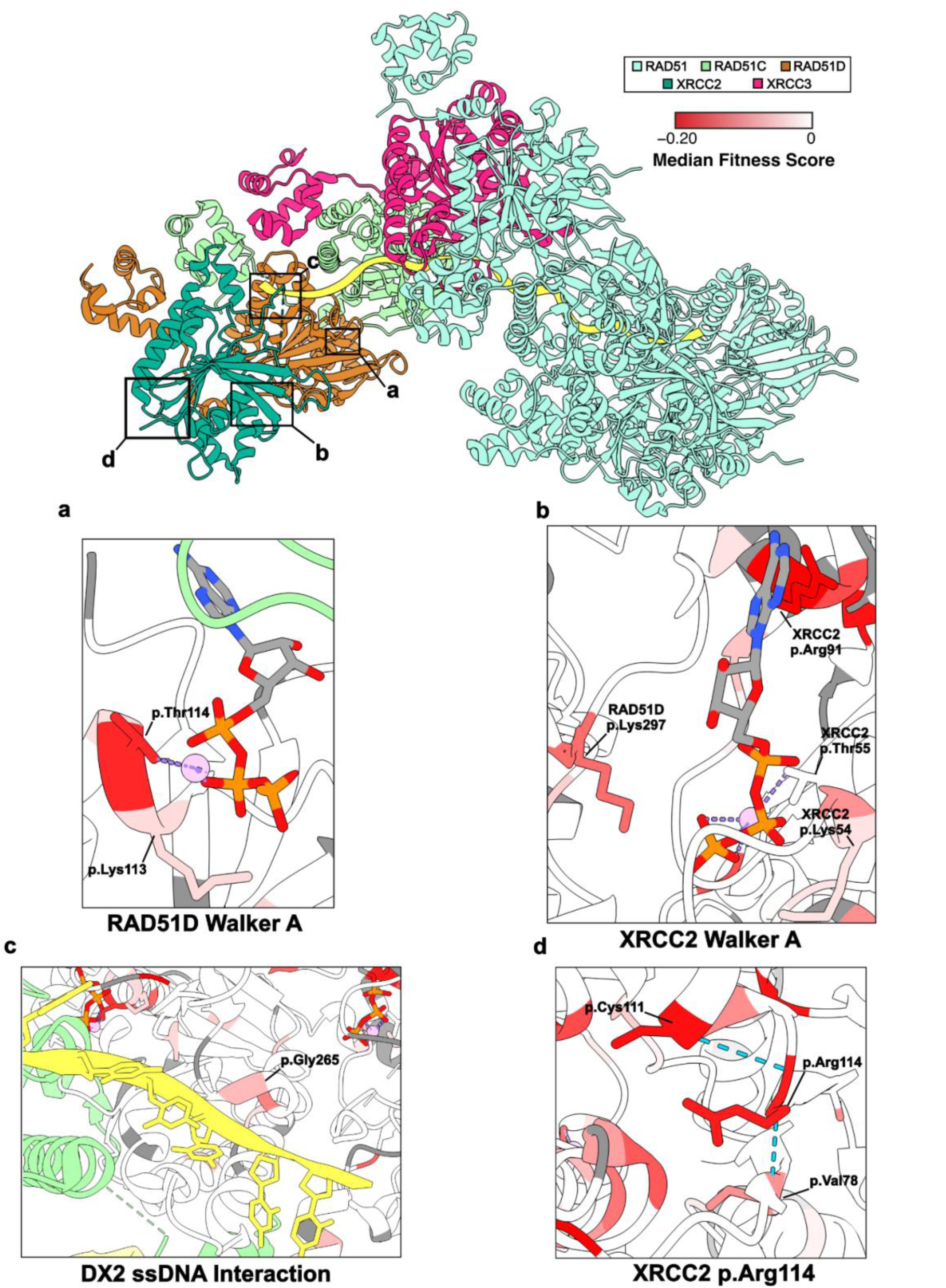
Structural consequences of RAD51D and XRCC2 missense variants on the X3CDX2 complex. (a) Diagram of RAD51D’s Walker A ATP-binding motif (PDB: 9SVX^26^) colored by median fitness score as indicated (gray denotes residues not scored). RAD51D p.Lys113 and p.Thr114 are highlighted. RAD51C is colored in green. (b) Diagram of XRCC2’s Walker A ATP-binding motif (PDB: 9SVX^26^) colored by median fitness score. RAD51D p.Lys297 and XRCC2 p.Lys54, p.Thr55, and p.Arg91 are highlighted. (c) ssDNA binding motif of RAD51D and XRCC2 (PDB: 9SVX^26^) colored by median fitness score. Bound ssDNA is colored in yellow. RAD51C is colored in green. RAD51D p.Gly265 is highlighted. (d) XRCC2 p.Arg114 (PDB: 9SVX^26^) and potential interacting partners colored by median fitness score. Likely hydrogen bonding partners colored by median fitness score. Dotted light blue lines highlight potential hydrogen bonding interactions.

**Extended Data Fig. 10.**
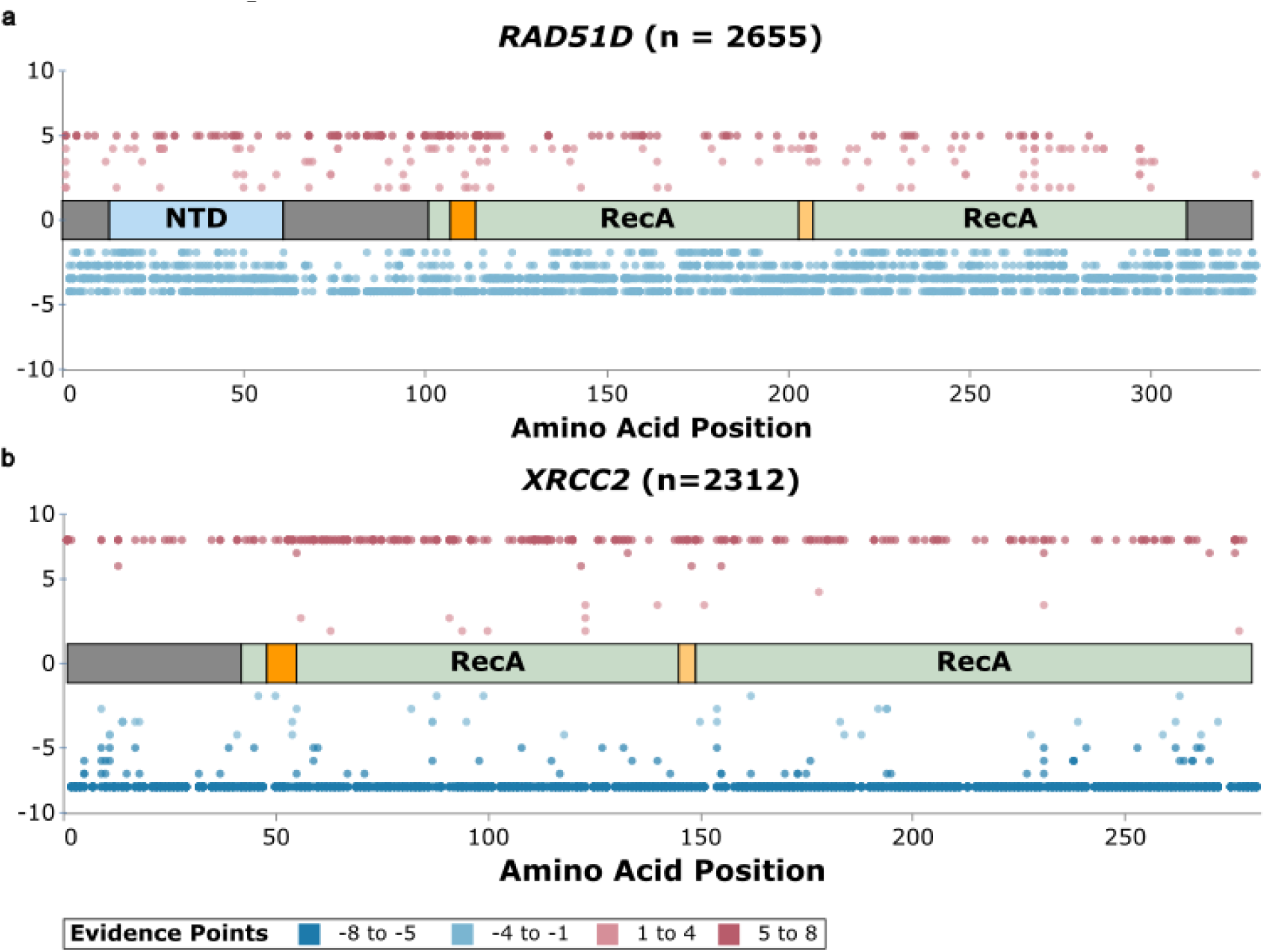
Calibrated ACMG/AMP evidence points for *RAD51D* and *XRCC2* variants mapped across protein domains. (a) Calibrated evidence points for *RAD51D* SNVs (n=2,655) plotted by amino acid position. Positive values (pink/red) indicate pathogenic evidence and negative values (blue) indicate benign evidence, with color intensity reflecting number of evidence points received. Protein domain architecture is shown along the midline: N-terminal domain (NTD, light blue), RecA-like domains 1 and 2 (green), Walker A and Walker B motifs (orange), and unannotated linker regions (grey). Variants in the C-terminal exon 10 region (residues 301–328) predominantly receive benign evidence. (b) As described in (a), for *XRCC2* SNVs (n=2,312). Domain architecture indicates the two RecA-like domains (green), Walker A and Walker B motifs (orange), and unannotated N-terminal region (grey).

